# MYB insufficiency disrupts proteostasis in hematopoietic stem cells leading to age-related neoplasia

**DOI:** 10.1101/2021.09.04.458970

**Authors:** M.L. Clarke, R.B. Lemma, D.S. Walton, G. Volpe, B. Noyvert, O.S. Gabrielsen, J. Frampton

## Abstract

The Myb transcription factor plays critical roles in normal and malignant hematopoiesis. Acquired genetic dysregulation of Myb, which plays a central role in hematopoietic stem cell (HSC) gene regulation, is involved in the etiology of a number of leukemias. Also, inherited non-coding variants of the Myb gene are a factor in susceptibility to many hematological conditions, including myeloproliferative neoplasms (MPN), but the mechanisms by which variations in Myb levels predispose to disease, including age-dependency in disease occurrence, are completely unknown. Here, we address these key points by showing that Myb insufficiency in mice leads in later life to MPN, myelodysplasia, and leukemia, mirroring the age profile of equivalent human diseases. This age-dependence is intrinsic to HSC, involving progressive accumulation of subtle changes. Interestingly, and linking to previous studies showing the importance of proteostasis to the maintenance of normal HSC, we observed altered proteosomal activity in young Myb-insufficient mice and later elevated ribosome activity. We propose that these alterations collectively cause an imbalance in proteostasis, potentially creating a cellular milieu favoring disease initiation by driver mutations.

## INTRODUCTION

The relationship between ageing and tissue stem cells is a complex interdependency that has considerable implications, not least for the occurrence of many chronic diseases. Both intrinsic and extrinsic aspects of the regulation of stem cell function are susceptible to the burden of changes accumulated with age, whether in the form, for example, of acquired DNA mutations or protein aggregates in the cell itself or an altered local or systemic environment including compositional variations in the ECM or the profile of inflammatory cells and mediators (Oh, Lee et al. 2014, Ermolaeva, Neri et al. 2018).

Hematological diseases such as myeloproliferative neoplasm (MPN) or myelodysplastic syndrome (MDS), which arise within the heterogenous population of hematopoietic stem cells (HSC) (Elias, Schinke et al. 2014, Mead and Mullally 2017), occur predominantly in later age, at least 60% of patients with blood malignancies being diagnosed over the age of 65. Multiple differences have been shown to characterise old versus young HSC (Sudo, Ema et al. 2000, Geiger, de Haan et al. 2013, Mejia-Ramirez and Florian 2020), including a shift to a more proliferative state and skewing of the balance of commitment to lineage-restricted progenitors. These functional deficiencies in HSC are paralleled by changes in the epigenome and transcriptome, including genes involved in inflammatory responses, as well as altered proteostasis and metabolism (Rossi, Bryder et al. 2005, Mejia-Ramirez and Florian 2020).

One or more of the intrinsic age-related changes in HSC must in some situations compound, or be compounded by, variations in the gene regulatory network that underpins their stem cell properties. Hence, GWAS studies have revealed that the age-dependent risk of MPN is linked with specific genes encoding epigenetic or transcriptional regulators. One such protein is the transcription factor Myb, which was shown to be linked to MPN in association with specific SNPs upstream of the gene that lead in a dominant fashion to lower levels of expression (Tapper, Jones et al. 2015). Myb has long been known to be crucial in regulating gene transcription in cells of the hematopoietic hierarchy, including in both progenitors and more mature cells, and to be a driver of some hematological malignancies and other types of cancer when its expression is dysregulated (Clappier, Cuccuini et al. 2007, Ramsay and Gonda 2008, Brayer, Frerich et al. 2016, Xu, Zhao et al. 2019)

Although lower Myb activity in model systems has negative impacts on HSC function (Sandberg, Sutton et al. 2005, Lieu and Reddy 2009), we observed that this can also confer stem cell-like properties on committed myeloid progenitors, producing an MPN-like condition (Garcia, Clarke et al. 2009, Clarke, Volpe et al. 2017). The latter study utilised a mouse line that exhibits a profound knockdown of the Myb protein, which is of the order of only 20% of normal levels (Emambokus, Vegiopoulos et al. 2003), and therefore is not very representative of the degree of variation that is expected within a human population as a consequence of the *Myb* gene upstream SNPs. Here, we have used a mouse model with a less pronounced reduction in Myb expression to explore how such changes might influence the occurrence of myeloid disease throughout the full lifespan. Our findings show that constitutive Myb haploinsufficiency leads to MPN / MDS at a high frequency in older animals, with features highly analogous to those seen in humans. Using a conditional haploinsufficiency strategy we further demonstrate that the level of Myb determines an age-dependent intracellular environment in respect to proteome homeostasis (‘proteostasis’), metabolic activity, and the rate of proliferation. We propose that these cellular perturbations either drive malignant transformation or provide conditions that favour clonal expansion following driver gene mutation. Focussing on disturbances in proteostasis related to Myb insufficiency, we highlight that these can be the result of a mixture of life-long effects, which are either directly or indirectly due to the action of Myb on specific target genes, and secondary effects that accumulate over time as a consequence of the primary effects.

## RESULTS

### Myb haploinsufficiency represents a model for transcription factor-dependent age-related development of MDS and MPN from HSC

Myb haploinsufficiency, achieved through heterozygosity for the constitutive knockout allele of the gene, has only relatively minor effects on the hematological status of young adults (Kasper, Boussouar et al. 2002), although the long-term consequences have not been determined. Therefore, as a potential model for Myb insufficiency in humans, cohorts of WT and *Myb*^+/−^ mice were aged for up to 22 months, monitoring blood values throughout for the development of disease (Supplementary Figure 1). Excluding animals that had to be sacrificed for reasons not related to their hematological status, the WT and *Myb*^+/−^ cohorts consisted of 19 and 17 animals, respectively. Disease was classified according to the Bethesda criteria for non-lymphoid hematopoietic neoplasms (Kogan, Ward et al. 2002).

Kaplan-Meyer survival plots for the two cohorts are shown in Figure 1A. All but one WT animal survived for 22 months with signs of hematological disease being seen in 4 animals (1 MDS, 2 MPN and 1 myeloid leukemia). In contrast, all but one animal in the *Myb*^+/−^ cohort developed myeloid disease with the percentage of survival at the end of the experiment being just 52% (log rank test, p = 0.0035). Eight of the *Myb*^+/−^ mice had to be sacrificed due to myeloid leukemia (comprising 50% monocytic, 38% MPN-like, and 13% myelomonocytic) and one because of a lethal myeloproliferation. The profile of non-fatal myeloid disease phenotypes amongst the *Myb*^+/−^ cohort included two MDS and 5 MPN (Figure 1B).

**Figure 1.**
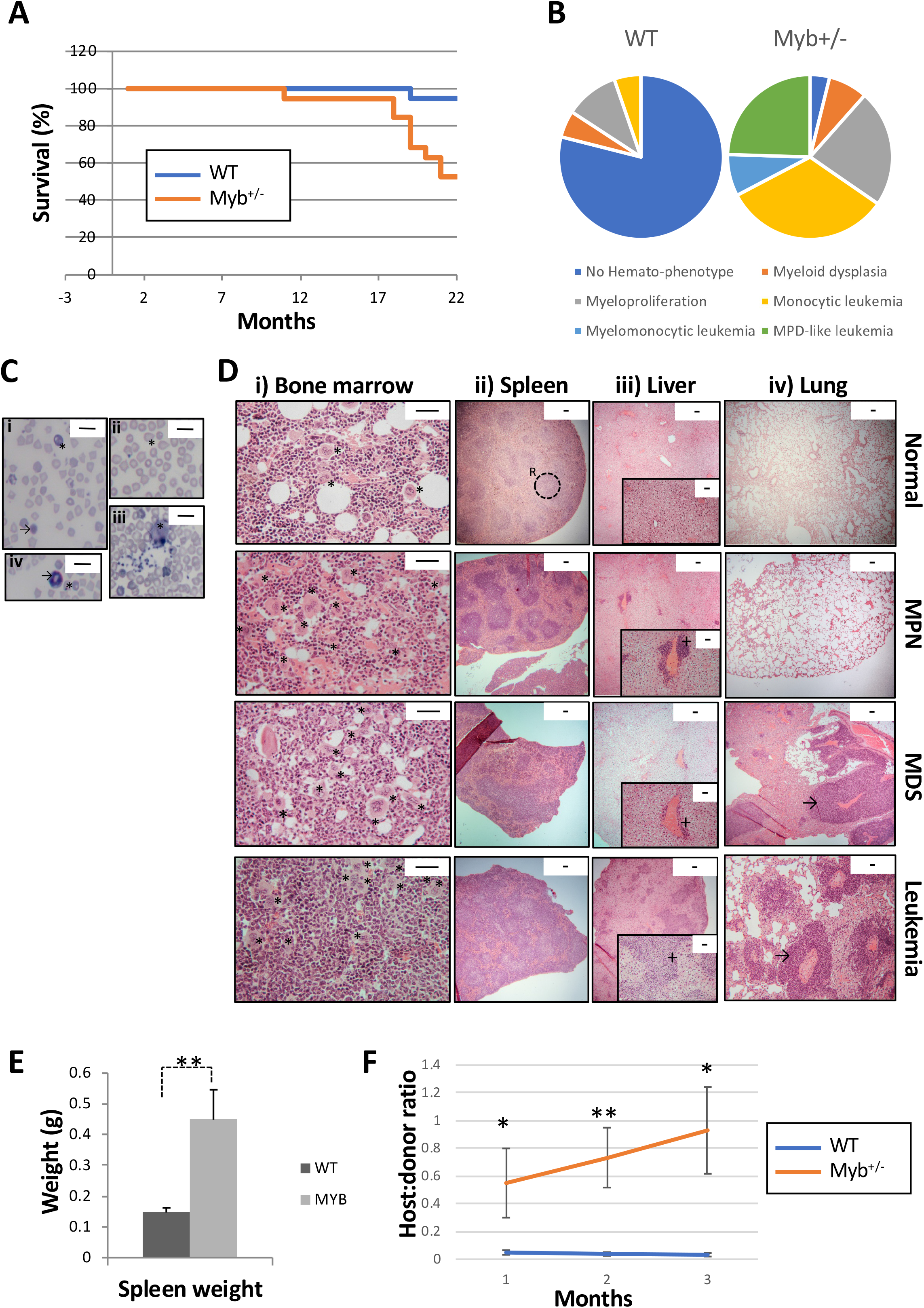
Myb deficiency leads to age-related hematological disease. A) Kalpan-Meier curve showing the percentage survival of a cohort of ageing WT and *Myb*^+/−^ mice (n=19 and 17 respectively) over 22 months. Significance was calculated using the log rank test with p = 0.0035. B) Pie charts depicting the types of myeloid disease arising with age. C) Blood smears from diseased mice showing examples of abnormal peripheral blood cells. i) Target cell erythrocyte (codocyte) (*). Polychromatic erythrocyte (→). ii) Teardrop erythrocyte (dacrocyte) (*). iii) Micromegakaryocte (*). iv) Howell-Jolly body in erythrocytes (*). Ring-neutrophil (→). Scale bar = 20μm. D) Representative sections from tibial bone, spleen, liver and lung from mice characterised as WT, MPN, MDS, or myeloid leukemia. Scale bars are indicated. i) Bone section with megakaryocytes (*) showing an increase in diseased mice. Scale bar = 50 μm. ii) Spleen sections showing disrupted red and white pulp structure in diseased mice. An example area of white (circle) and red (R) pulp is indicated. Scale bar = 100 μm. iii) Liver sections showing infiltration of myeloid cells (+). Scale bar = 100 μm (inset: scale bar = 50μm) iv) Lung sections showing infiltration of myeloid cells (→). Scale bar = 100 μm. E) Spleen weight (g) of ageing WT and *Myb*^+/−^ mice at the point of culling (n=19 and 17, respectively). Significance was calculated using Student’s t-test p=0.006. Data is represented as the mean and SEM. F) Ratio of host:donor bone marrow cells from WT (n=4) and mice with myeloid leukemia (n=6). Whole bone marrow was transplanted into sub-lethally irradiated (450Gy) B6.SJL-*Ptprc*^*a*^ / BoyJ mice. Ratios are calculated by antibody staining of peripheral blood against CD45.1 and CD45.2 antigens at 1-, 2- and 3-months post-transplant (p=0.043, 0.007 and 0.011, respectively). Data points represent the mean ratios with SEM.

The MPN phenotypes arising in *Myb*^+/−^ animals were characterised in peripheral blood by thrombocytosis and leucocytosis, whereas those mice with MDS exhibited combinations of thrombocytopenia, neutropenia, lymphopenia, and anemia (Supplementary Figure 2). Peripheral blood smears from mice with MDS phenotypes revealed significant dyserythropoiesis, including polychromasia, macrocytes, target cells, Howell-Jolly bodies, dacrocytes (tear-drop cells) and helmet cells, together with the presence of micromegakaryocytes in the periphery (Figure 1C). Neutrophil dysplasia was also observed in the form of excess immature band neutrophils and hyper-segmented granulocytes (Figure 1C).

Bone sections (Figure 1D) from *Myb*^+/−^ mice that developed MPN revealed hypercellular bone marrow with increased megakaryocytes (asterisks), which typically occurred in clusters and possessed characteristic stag-horn nuclei. MDS was distinguishable by increased cellularity and the presence of dysplastic megakaryocytes such as micromegakaryocytes and hypolobulated megakaryocytes.

Splenomegaly was observed in the majority of *Myb*^+/−^ diseased mice (Figure 1E) and sections of spleen from mice exhibiting MPN demonstrated enlarged red pulp indicative of extramedullary hematopoiesis. Minor hepatic infiltration was also observed, but no lung involvement (Figure 1D). Analysis of peripheral organ sections from mice exhibiting MDS revealed highly disrupted splenic architecture with the presence of extramedullary megakaryocytes, as well as hepatic and lung infiltration (Figure 1D).

*Myb*^+/−^ mice that developed myeloid leukemia possessed features of either MPN or MDS suggesting a progression to leukemia from these phenotypes. Tissue sections showed a marked increase in megakaryocytes in both the bone marrow and spleen, which had a highly disrupted architecture. The liver and lung exhibited extensive infiltration of hematopoietic cells (Figure 1D). Leukemia progression was confirmed in these mice by transplantation of whole bone marrow cells into sub-lethally irradiated recipients followed by peripheral blood analysis of engraftment. Compared to cells from WT mice, the leukemic *Myb*^+/−^ bone marrow engrafted at a high rate and transferred the phenotype to the recipient mice (Figure 1F).

### The age dependence of Myb insufficiency-associated disease is largely an intrinsic feature of HSC

The majority of animals in the *Myb*^+/−^ cohort exhibited a hematological phenotype beyond one year of age (average 13.8 months), which might be the result of Myb-dependent mechanisms solely intrinsic to HSC / progenitor cells or alternatively could be a consequence of an interaction between the lower levels of the transcription factor and age-related changes in the environment the cells experience. To try to discriminate between these possibilities, use was made of mice in which a *Myb* allele can be conditionally ablated (Emambokus, Vegiopoulos et al. 2003) together with a Tamoxifen-inducible Cre recombinase (CreER^T2^ (Metzger, Clifford et al. 1995)) and a reporter transgene (*mTmG*, (Muzumdar, Tasic et al. 2007)) that can be switched from expression of tdTomato to Green Fluorescent Protein (GFP) following Cre-mediated recombination. The strategy, summarised in Figure 2A, entailed inducing deletion of one allele of *Myb* in animals at 2 or 12 months of age followed by tracking of the incidence of myeloid disease features. Cohort 1 consisted of 10 *Myb*^+/+^:*CreER*^*T2*^:*mTmG* and 11 *Myb*^+/*F*^:*CreER*^*T2*^:*mTmG* mice that were treated with Tamoxifen at 2 months of age, while Cohort 2 comprised 10 *WT*:*CreER*^*T2*^:*mTmG* and 10 *Myb*^+/*F*^:*CreER*^*T2*^:*mTmG* mice in which *Myb* allele deletion was induced at 12 months of age. The inclusion of the *mTmG* transgene reporter enabled assessment of the effectiveness of the deletion and identification and sorting of cells that either deleted (GFP^+^) or failed to delete (tdTomato^+^). The efficiency of deletion (~50%) was confirmed by quantitative PCR of the relevant genomic interval using DNA prepared from whole peripheral blood cells following red cell lysis (Figure 2B). A cohort of *Myb*^+/+^:*CreER*^*T2*^:*mTmG* animals was used at both time points to control for Tamoxifen-specific effects. A small cohort of *Myb*^+/−^ mice were also injected with Tamoxifen to demonstrate that the agonist does not accelerate the hematopoietic phenotype previously observed, 80% of the *Myb*^+/−^ mice survived up to 18 months of age (the average onset of disease in aged *Myb*^+/−^ mice) (Supplementary Figure 3A).

**Figure 2.**
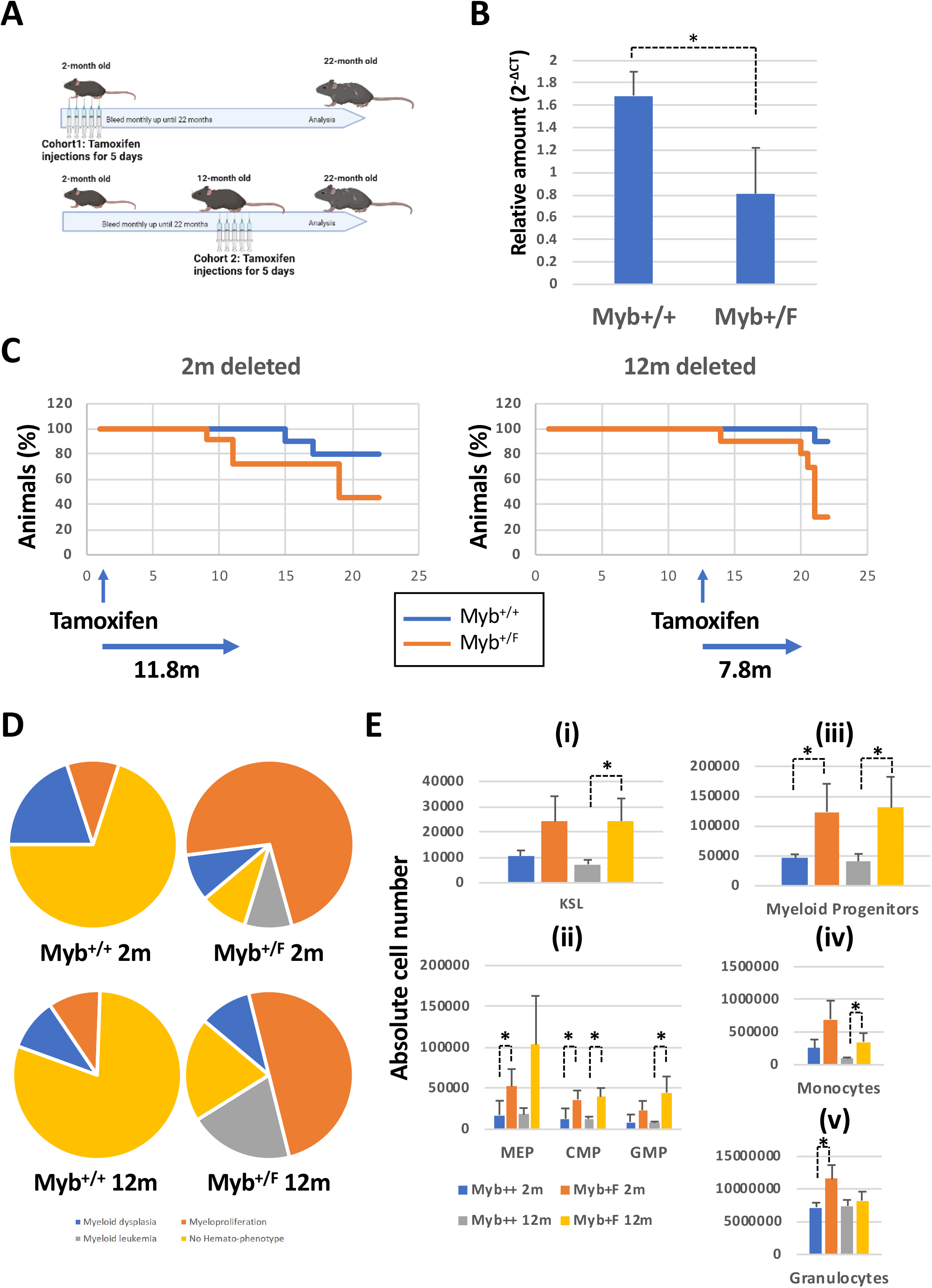
Conditional deletion of Myb at 2- or 12-months determines disease is HSC intrinsic. A) Schematic depicting the Tamoxifen treatment of *Myb*^+/+^:*CreER*^*T2*^:*mTmG* (*Myb*^+/+^) and *Myb*^+/*F*^:*CreER*^*T2*^:*mTmG* (*Myb*^+/*F*^) mice at either at 2- or 12-months of age. B) Quantitative PCR on peripheral blood genomic DNA, detecting the relative levels of *Myb*^*ex5*^ product relative to an internal control *GpIIb* calculated using the ΔCT method. Values represent the mean expression plotted as 2^−ΔCT^ with SEM (n=10 p=0.020). C) Kalpan-Meier curve of the survival over a period of 22 months of *Myb*^+/+^ and *Myb*^+/*F*^ mice after Myb deletion at 2- or 12-months. Arrows signify the average time from tamoxifen deletion until disease appearance. Significance was calculated using the log rank test with p = 0.026 and 0.006 for the 2- and 12-month deleted, respectively. n=10 for *Myb*^+/+^ 2- and 12- months, n=11 for *Myb*^+/*F*^ 2-months, n=10 for *Myb*^+/*F*^ 12-months. D) Classification of myeloid disease arising in the ageing cohorts of Myb deleted mice. E) Absolute bone marrow cell numbers based on antibody staining and total bone marrow count. KSL: Kit^+^Sca^+^Lin^−^; MEP: Megakaryocyte-Erythroid Progenitor; CMP: Common Myeloid Progenitor; GMP: Granulocyte-Monocyte Progenitor; Myeloid progenitors: Kit^+^Sca^−^Lin^−^; Monocytes: CD11b^+^Gr1^−^; Granulocytes: CD11b^+^Gr1^+^. n=10 for *Myb*^+/+^ 2- and 12-months, n=11 for *Myb*^+/*F*^ 2-months, n=10 for *Myb*^+/*F*^ 2-months. Significance was calculated by Student’s T-test. i) KSL^12m^ p=0.032. ii) MEP^2m^ p=0.052, CMP^2m^ p =0.023, CMP^12m^ p=0.014, GMP^12m^ p=0.031. iii) Prog^2m^ p=0.052, Prog^12m^ p=0.047. iv) Monocytes^12m^ p= 0.028, v) Granulocytes^2m^ p= 0.032.

As illustrated in the Kaplan-Meyer graphs (Figure 2C), the deletion of one *Myb* allele at either 2 or 12 months caused a statistically significant reduction in the survival of *Myb*^+/*F*^:*CreER*^*T2*^:*mTmG* mice compared to the control group (log rank test: p = 0.026 and p = 0.0062, respectively). As for the aged constitutive *Myb*^+/−^ animals, not all of the mice subjected to conditional deletion of *Myb* succumbed to disease, however at the end of the experiment at 21 months it was obvious that a significant proportion had developed myeloid disease. In cohort 1, ten of the *Myb*^+/*F*^:*CreER*^*T2*^:*mTmG* mice showed signs of disease (1 MDS, 8 MPN, 1 myeloid leukemia) compared to just two of the *Myb*^+/+^:*CreER*^*T2*^:*mTmG* (1 MDS, 1 MPN). Eight of the Cohort 2 *Myb*^+/*F*^:*CreER*^*T2*^:*mTmG* mice developed myeloid disease, with a slightly higher incidence of myeloid leukemia compared to the 2-month deleted animals (20% and 9%, respectively) along with 1 MDS and 5 MPN (Figure 2D). Tissue sections were collected from the aged mice at the time of death and the features described earlier for the aged *Myb*^+/−^ mice were consistent here, enabling the determination of disease phenotype (Supplementary Figure 3B).

Analysis of the bone marrow from animals in both cohorts revealed that deletion of *Myb* at both 2- and 12-months of age led to an increase in the absolute number of HSC (KSL) and Kit^+^ myeloid progenitors (Figure 2E). Breaking down the progenitors by their lineage potential, there was an indication that the increase in GMP following *Myb* allele deletion was somewhat greater when this occurred at 12 months, being paralleled by an increase in monocytes within the bone marrow.

Given that a period of several months elapsed between induction of *Myb* allele deletion and the onset of signs of hematological disease, whether deletion was performed in 2- or 12- month old animals (average time delay 11.8 and 7.8 months, respectively), it seems that the age-related occurrence of MDS, MPN, or leukemia in *Myb*^+/−^ animals is to a large extent due to an accumulated consequence of the absence of Myb rather than non Myb-related ageing effects.

### Myb-insufficient HSC are functionally compromised

Since both MPN and MDS are thought to arise as a result of acquired mutations within the HSC compartment, we compared the profile and function of stem cells from young and aged WT and *Myb*^+/−^ mice. Blood values from 2-month old mice revealed that the *Myb*^+/−^ mice exhibit no distinct hematological pathology (Supplementary Figure 4), whereas in the bone marrow increased numbers of KSL HSC were observed (p=0.0009), including long-term HSC (KSL CD48^−^CD150^+^) (p = 0.013) (Figure 3A). By 12 months of age, although still exhibiting increased numbers of KSL HSC (not statistically significant), the relative number of long-term HSC was reduced (p = 0.005).

**Figure 3.**
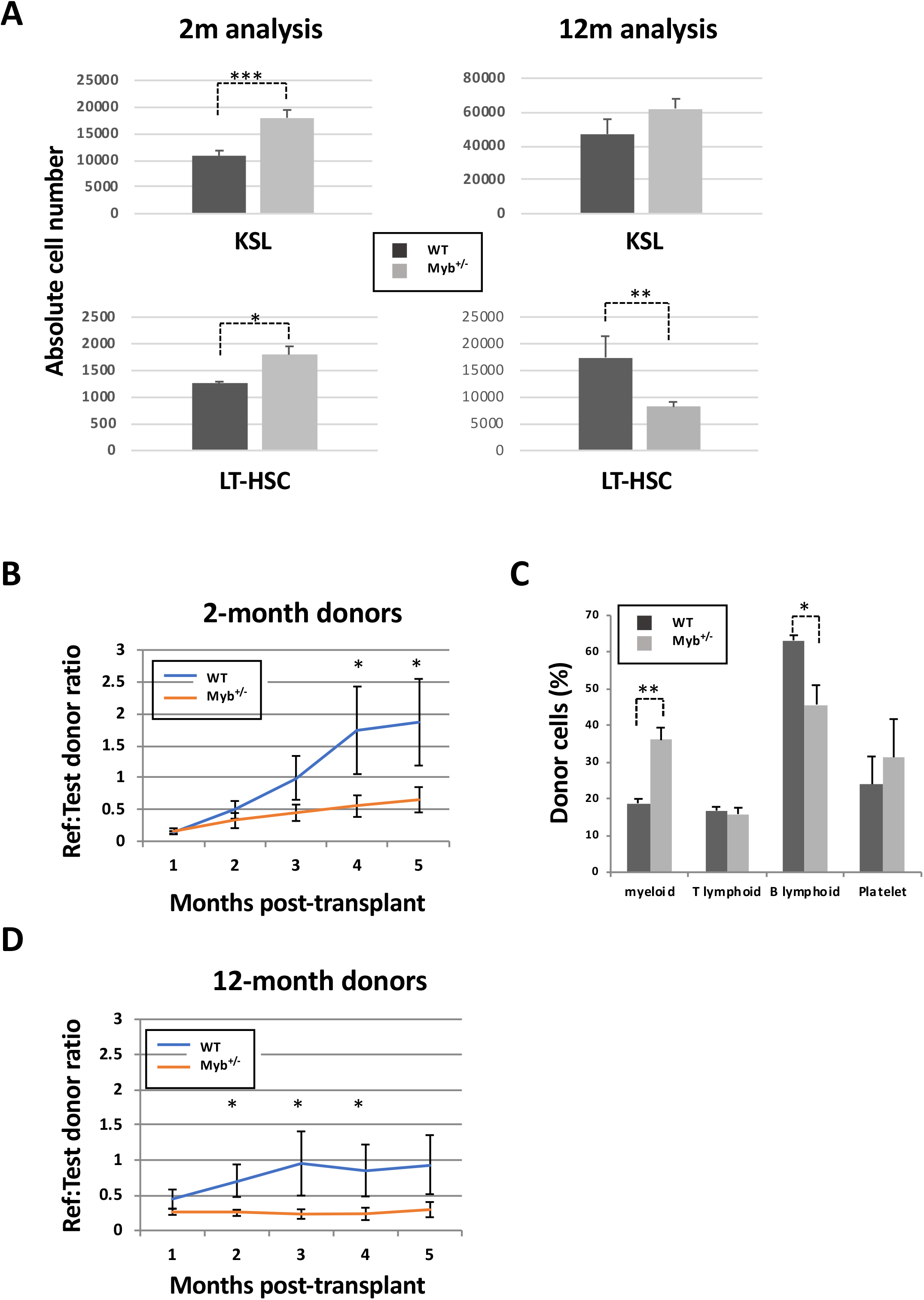
Myb deficient HSC are compromised in their function. A) Bone marrow absolute counts of KSL and LT-HSC (KSLCD48-CD150+) from 2- (n=8) and 12-month (n=8) old WT and *Myb*^+/−^ mice. KSL^2m^ p= 0.0009, KSL^12m^ p=0.053, LT-HSC^2m^ p=0.013, LT-HSC^12m^ p=0.005. B) Sorted KSL cells from 2- (n=9) and 12-month (n=9) old WT or *Myb*^+/−^ donors (carrying *vWF-tdTomato* transgene) plus reference bone marrow were transplanted into lethally irradiated recipients (900Gy). Reference:donor cell ratios were calculated by flow cytometry. P^2m^= 0.040, 0.038 and P^12m^=0.030, 0.006, 0.055. C) Analysis of donor cell percentage in the periphery of recipient mice (n=3) 4-months post-transplant receiving 2-month old KSL. Antibody staining gated on CD45.2 donor cells: Myeloid (CD11b^+^), T-lymphoid (CD4^+^CD8a^+^), B-lymphoid (B220^+^) and platelet (vWF^+^). Myeloid p= 0.001, B-lymphoid p=0.011)

In order to assess the effect of Myb insufficiency on stem cell function, including self-renewal and lineage commitment potentials, transplantation assays were performed on sorted KSL HSC from 2-month old WT and *Myb*^+/−^ mice containing the *vWF*-<u>*tdtomato*</u> transgene (Sanjuan-Pla, Macaulay et al. 2013) in order to facilitate tracking of platelet differentiation that would otherwise not be detected by immunofluorescent staining for CD45. Engraftment rates over a 5-month period revealed that *Myb*^+/−^ HSC are significantly impaired in their ability to reconstitute the hematopoietic system of lethally irradiated recipients (P < 0.05) (Figure 3B). Analysis of the donor-derived cells within the peripheral blood showed that the *Myb*^+/−^ HSC give rise to a higher proportion of myeloid cells (CD11b^+^) (P = 0.001) and that the reduced engraftment is not an artefact of increased platelet numbers, since the CD45^-^ vWF^+^ fraction of the peripheral blood is not different for WT and *Myb*^+/−^ donor cells (Figure 3C). A similar difference was observed using HSC from 12-month old WT mice although the significant reduction in engraftment was evident much earlier following transplantation compared to the 2-month old donor cells (2-versus 4-months respectively) (Figure 3D). As expected the 12-month old WT KSL cells engrafted at a lower efficiency compared to younger WT KSL cells.

### Distinct gene expression changes resulting from Myb insufficiency are seen when comparing HSC from young versus old mice

Further to assess how HSC function might be affected by *Myb* deficiency, RNA-Seq transcriptome analysis was performed on sorted KSL HSC derived from 2- and 12-month-old WT and *Myb*^+/−^ mice. Biological triplicates were performed for each of the four categories. Considering expression differences with an adjusted-p value of less than 0.1, 247, 1108, 188, and 1433 genes were identified as being differentially expressed comparing *WT* 2- versus 12-month old, *Myb*^+/−^ 2- versus 12-month old, 2-month old *WT* versus *Myb*^+/−^, and 12-month old *WT* versus *Myb*^+/−^, respectively. The overlap of genes uniquely shared as differences in 2-month old *Myb*^+/−^ and aged *WT* HSC was just 4 (2.1% of the 2-month old *WT* versus *Myb*^+/−^ differences, and 1.6% of the *WT* ageing differences) (Figure 4A), implying that young *Myb*^+/−^ HSC are not particularly prone to a premature ageing phenotype, but instead exhibit other biological process alterations that affect HSC function and ultimately renders them susceptible to neoplastic disease.

**Figure 4.**
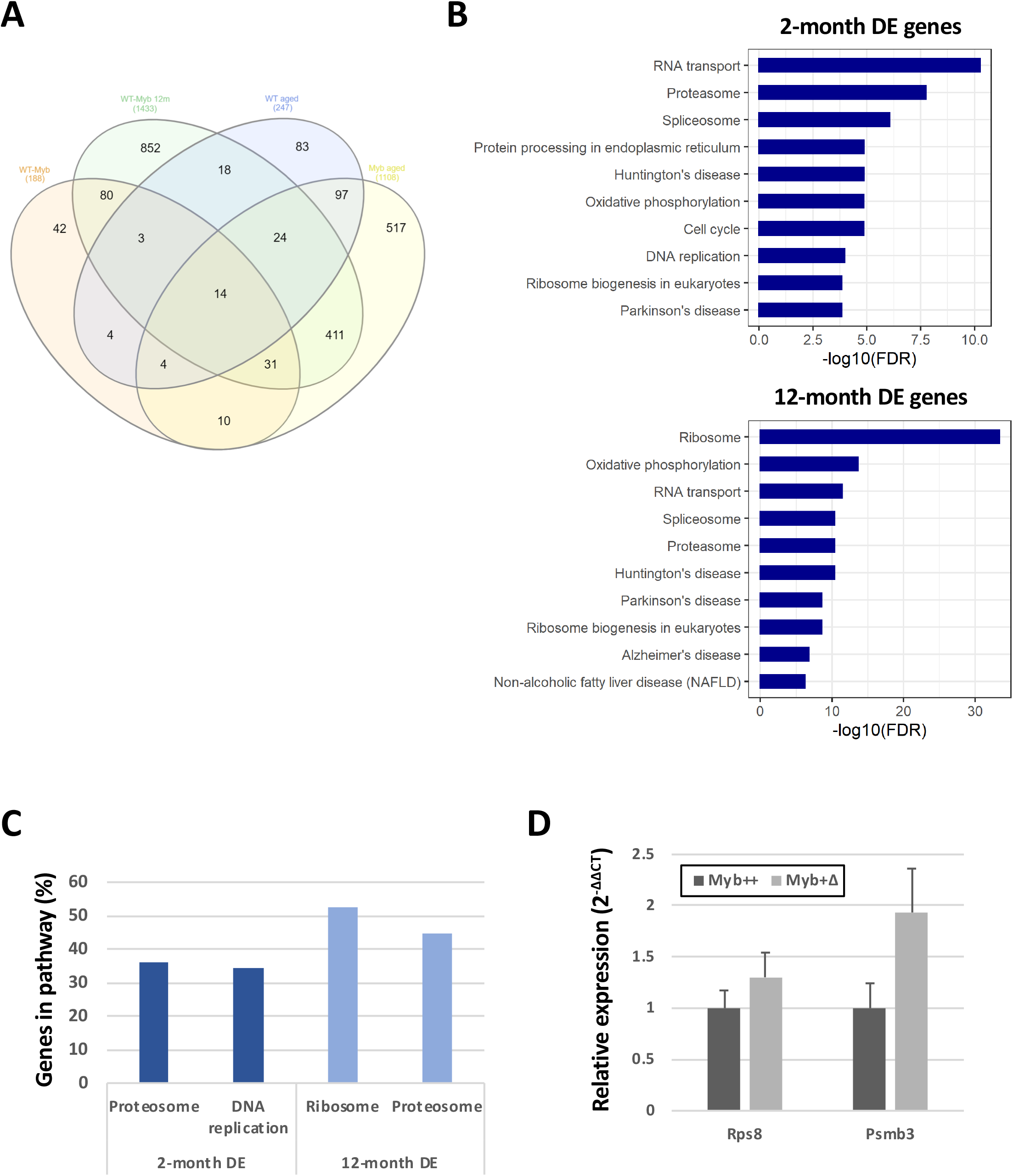
RNA-Seq analysis determines key pathways that are altered by low levels of Myb. A) Venn diagram of the significantly (adjusted p=0.1) differentially expressed (DE) genes from RNA-Seq comparisons of WT and *Myb*^+/−^ KSL young (2-month) versus aged (12-month). The numbers of genes overlapping are indicated. B) KEGG pathway analysis from the comparison of 2-month WT vs *Myb*^+/−^ and 12-month old WT and *Myb*^+/−^ KSL cells. The top 10 most significant pathways are depicted based on their FDR. C) The percentage of genes in each depicted pathway that are significantly DE in 2- and 12-month old KSL. D) Quantitative PCR of RNA expression in aged *Myb*^+/+^:*CreER*^*T2*^:*mTmG* and *Myb*^+/Δ^:*CreER*^*T2*^:*mTmG* sorted GFP+ KSL cells. Values represent the mean expression plotted as 2^-ΔΔCT^ with SEM n=3.

Analysis of the enriched KEGG pathways in the comparison between 2-month old *WT* versus *Myb*^+/−^ revealed an enrichment in genes encoding proteins associated with the proteosome, which were similarly predominant in the differences distinguishing 12-month old *WT* from *Myb*^+/−^ HSC (Figure 4B,C and Supplementary Figure 5). *Myb*^+/−^ HSC at 2 and 12 months of age showed higher expression of 17 and 33 proteosome-associated genes, respectively, encoding components of the 19S and 20S proteasome complexes that together constitute the 26S proteasome (Figure 4B), which is crucial for the normal functioning of HSC (Hidalgo San Jose, Sunshine et al. 2020). In addition to enrichment of proteasome-associated genes, the 12-month old *Myb*^+/−^ HSC showed highly significant enrichment of genes encoding proteins linked with the ribosome (Figure 4B,C); these changes were only seen at this age and relate to rRNA processing and ribosome biogenesis. A total of 93 genes encoding ribosomal components and 34 genes involved in ribosomal biogenesis are increased in the *Myb*^+/−^ HSC at 12 months of age (Supplementary Figure 6). Another interesting pathway enrichment characterising 2-month old *Myb*^+/−^ HSC involves genes connected to DNA replication and cell cycle, including components of the DNA polymerases and MCM complexes (Figure 4B,C and Supplementary Figure 7). As for the proteasome component differences, the same DNA replication associated genes are also seen to be enriched in 12-month old *Myb*^+/−^ HSC compared to their WT equivalents at this age.

HSC gene expression differences were also compared between aged WT and *Myb*^+/*F*^ mice rendered haploinsufficient (*Myb*^+/*Δ*^) by Cre-mediated deletion at 2-months of age. HSC in which deletion had occurred, identified on the basis of their expression of GFP from the recombined mTmG reporter, were sorted and quantitative RT-PCR performed on representative genes highlighted in the differential transcriptome comparison of *WT* and constitutively *Myb*^+/−^ mice. HSC were collected from animals at 14-months of age, that is at the average point of disease incidence following *Myb* allele deletion. *Myb*^+/*Δ*^ HSC showed a higher expression of genes associated with the ribosome (*Rps8*) and the proteosome (*Psmb3*), consistent with the observations seen in the aged cohort of *Myb*^+/−^ mice (Figure 4D).

### Changes in proteostasis-related gene expression associated with Myb-insufficiency and ageing is reflected in functional process differences in HSC

Based on the transcriptome analyses, it appears that groups of genes reflecting major cellular processes are affected by Myb insufficiency in HSC, either irrespective of age or as a compound effect with ageing. In the former category, genes associated with the proteasome and DNA replication, while changes in genes linked to ribosome structure and biogenesis become more obvious in older Myb insufficient animals. In order to confirm that the observed changes in gene expression are reflected in alterations in the corresponding cellular processes, assays were performed that relate to the collective activity of the relevant encoded proteins in relation to proteostasis (Figure 5) and DNA replication (Supplementary Figure 8).

**Figure 5.**
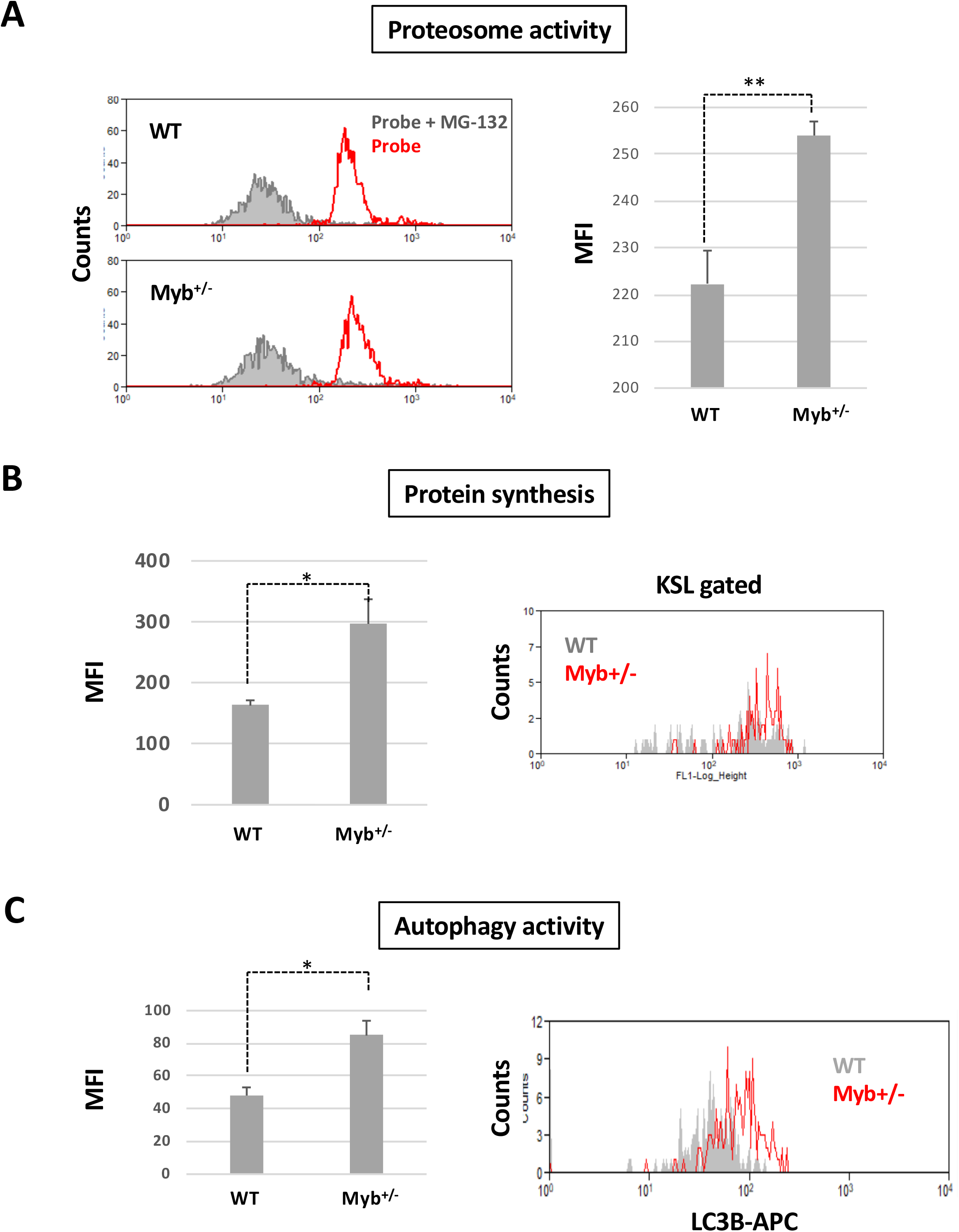
Myb deficiency affects protein production and degradation. A) KSL cells from 2-month old WT and *Myb*^+/−^ mice were stained with Me4BodipyFL-Ahx3Leu3VS proteosome activity probe either with or without prior incubation with MG-132. Median fluorescence intensity (MFI) was calculated. n=3, p= 0.007. B) Protein synthesis was assessed in 12-month old KSL cells from WT and *Myb*^+/−^ mice using the OPP assay. Median fluorescence intensity (MFI) was calculated. n=3, p= 0.016. C) Staining against LC3B for detection of autophagy was performed in 2-month old KSL from WT and *Myb*^+/−^ mice. MFI was calculated. n=2, p= 0.025.

The activity of the proteasome was assessed in 2-month old KSL HSC isolated from both WT and *Myb*^+/−^ mice. Sorted HSC were stained for one hour using the Me4bodipyFL-Ahx3Leu3VS proteosomal activity probe, either with or without pre-treatment with the proteosome inhibitor MG132, and were then analysed by flow cytometry (Figure 5A). Median fluorescent intensity (MFI) analysis demonstrated a significant 15% higher proteasome activity (p = 0.007) in the *Myb*^+/−^ HSC.

Similarly, to determine whether the increase in ribosome-associated gene expression in 12-month old *Myb*^+/−^ HSC is reflected in an overall increase in protein synthesis, newly translated proteins were labelled using O-propargyl-puromycin (OPP) together with Click-iT technology and analysed by flow cytometry (Figure 5B). 12-month old *Myb*^+/−^ HSC showed a significant 2-fold higher MFI (p = 0.016) compared to equivalent WT cells, indicating a significant increase in protein synthesis in the aged Myb-insufficient stem cells.

An alternative protective mechanism the cell utilises to clear damaged cellular components and provide extra nutrients in times of proliferative expansion and stress is autophagy. Since we have observed an increase in proteosome activity concurrent with an increase in DNA replication genes, we wanted to assess whether the *Myb*^+/−^ cells exhibit an increase in autophagy. Using intracellular antibody staining against the autophagosome marker LC3B, we identified a significant (p = 0.025) increase in MFI in KSL from *Myb*^+/−^ compared to WT bone marrow (Figure 5C).

Overall, the functional assays show that the alterations in proteostasis- and DNA replication-associated gene expression as a consequence of Myb insufficiency have a measurable impact on the corresponding biological processes.

### A subset of genes encoding proteostasis-associated proteins are regulated by Myb in both mouse HSC and a human HSPC line

In order to confirm that the observed changes in gene expression in *Myb*^+/−^ mice are functionally conserved and to explore if the genes affected by Myb insufficiency are direct target genes of the transcription factor, we took advantage of recent transcriptome and ChIP-seq data for MYB from human K562 cells (Fuglerud, Lemma et al. 2017, Lemma, Ledsaak et al. 2021). K562 is a human chronic myeloid leukemia (CML) cell line and a model of HSPC. A total of 525 genes in K562 were defined as direct target genes of MYB based on their response to MYB knock-down and rescue, and their association with one or several MYB ChIP-peaks using the STITCHIT algorithm (Schmidt et al., 2019). The corresponding mouse genes (n=523) were identified as described in Materials and Methods. Using this gene set, corresponding to human direct target genes of MYB, we investigated its overlap with the significantly differentially expressed mouse genes analysed above but with a stringent filter (p < 0.05, Log2FC ≥ 0.5, n=662). This overlap analysis resulted in n=32 genes.

Enrichment analysis was performed on this gene set (n=32) using the *clusterProfiler* R package (version 3.12.0) (Yu, Wang et al. 2012). The C2 curated list of genes from MSigDB was utilized to investigate relevant pathway and functional terms. This analysis revealed top significant enrichments of proteasome- and transcription initiation-/RNA-polymerase-associated terms both in Wiki and KEGG pathway databases (Figure 6A). Among the 32 genes, 8 were proteasome component-encoding (*PSM*) genes. To further support the direct functional link between the top ranked proteasome-associated genes and MYB, we visualized the direct MYB ChIP-seq peak occupancy at the loci of selected *PSM* genes (Figure 6B, left panels). Moreover, we took advantage of an RNA-seq dataset from K562 cells (Fuglerud et al. 2017) and added the expression profiles of these selected genes in K562 cells upon MYB knockdown and rescue. Consistently, the expression of the selected proteasome-associated genes showed a significant increase when MYB was knocked down (Figure 6B, right panels). This is true for all the eight *PSM* genes (not shown).

**Figure 6.**
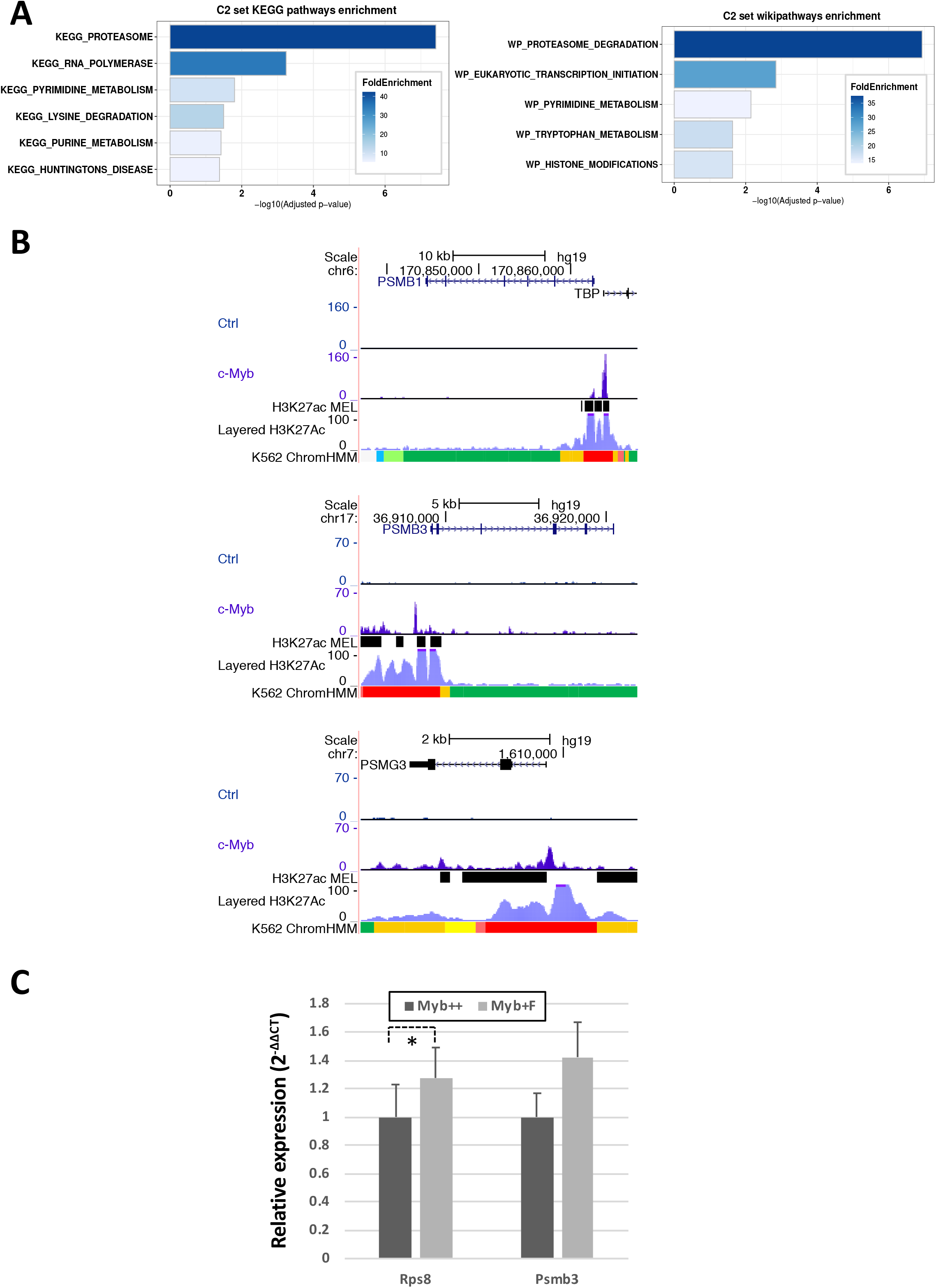
A subset of genes encoding proteostasis-associated proteins are directly regulated by Myb in both mouse HSC and a human HSPC line. A) Functional enrichment of the subset of mouse Myb insufficiency-responsive genes where their human homologues are direct target genes of MYB. Results from Wikipathways (upper panel) and KEGG pathway (lower panel) are shown. B) A subset of Myb insufficiency-responsive genes are direct target genes of MYB in K562 cells. Left panels: UCSC tracks showing MYB occupancy at the *PSMB1*, *PSMB3* and *PSMG3*. In addition to the K562 ChIP-seq tracks we incorporated H3K27Ac tracks (Accession ID: ENCFF117MIF) from the MEL mouse cell line, which is described as analogous to K562 cells in the ENCODE database. To make the MEL cell line derived H3K27Ac ChIP-seq peaks compatible with human hg19 coordinates, we used the UCSC *liftOver* tool (Hinrichs, 2006) before uploading it to the MYB UCSC session and visualizing the tracks using the UCSC genome browser (Kent et al., 2002). Right panels: expression profiles of *PSMB1*, *PSMB3* and *PSMG3*, respectively in K562 cells after siRNA KD of MYB and rescue by transient expression of 3✕TY1-MYB. C) Quantitative PCR of RNA expression in *Myb*^+/+^:*CreER*^*T2*^:*mTmG* and *Myb*^+/*F*^:*CreER*^*T2*^:*mTmG* sorted GFP+ KSL cells 24 hours post-deletion. Values represent the mean expression plotted as 2^−ΔCT^ with SEM. n=4 p= 0.047.

In order to confirm that Myb regulates the expression of mouse proteosomal genes in a similar manner to what was observed for the human genes, we performed conditional deletion on *Myb*^+/+^:*CreER*^*T2*^:*mTmG* and *Myb*^+/*F*^:*CreER*^*T2*^:*mTmG* mice by intraperitoneal injection of Tamoxifen. 24 hours later KSL HSC that had transitioned from tomato-FP to GFP were sorted and quantitative RT-PCR performed to measure expression of *Psmb3* RNA. Expression of Rps8 RNA was used as a control to determine if there is any change in ribosomal genes. *Myb*^+/*F*^:*CreER*^*T2*^:*mTmG* mice showed significantly higher expression of *Psmb3* (p = 0.047) with no significant change in in the expression of the ribosomal gene *Rps8* (Figure 6C).

## DISCUSSION

In any differentiation hierarchy that underpins tissue homeostasis, the transcriptional regulation of gene expression plays a major part in controlling stem cell maintenance and commitment, and maturation of the functional cell lineages. It is clear, especially from studies of hematopoiesis, that networks involving many transcription factors provide a mechanism for the control of gene expression that can sensitively respond to external signals reflecting changes in the tissue and its requirements for differentiated cells (Gottgens 2015). Such networks encompass multiple cross-regulatory interactions as well as positive and negative feedback loops. Any single alteration in an individual transcription factor, whether acquired through gene mutation or epigenetic modification, or inherited as a natural variation, might be expected to disturb the network balance. Although such a disturbance may only be subtle or potentially mitigated through the inherent compensatory capacity in the network, the consequence may nevertheless be a significant change in cell function, which could result in a disease initiation or facilitation (Assi, Bonifer et al. 2019).

The Myb transcription factor is known to play critical roles at all levels of the hematopoietic hierarchy, perturbations in its level of activity leading to cell deficits, cellular dysfunction, and malignancy (Ramsay and Gonda 2008, Wang, Angelis et al. 2018). Our previous studies, largely using mouse models of Myb deficiency, have focussed on the consequences in HSPC and illustrate that the requirement for a precise level of Myb activity is cell type dependent and that both loss- and gain-of-function can result when the correct level is disturbed. Hence, in the context of a Myb knockdown mouse model, HSC function is severely compromised, while at the same time downstream committed myeloid progenitors acquire stem cell-like characteristics, including self-renewal potential (Garcia, Clarke et al. 2009, Clarke, Volpe et al. 2017). A natural inherited variation in the regulation of *Myb* gene expression, involving upstream non-coding sequences, has been found to be associated with a wide range of hematological consequences, including MPN (Thein, Menzel et al. 2007, Tapper, Jones et al. 2015). In the latter instance, the risk-associated variants were shown to lower expression of Myb in HSPC throughout life, but it is unclear how this is related to disease occurrence in older age.

Mice that are haploinsufficient for Myb expression by virtue of one allele being knocked out are in most respects hematologically indistinguishable from their wild type counterparts, at least as young to middle aged adults (up to one year of age), although more profound Myb deficiency scenarios, either resulting from a majority knockdown or an activity-limiting mutation, are associated with significant perturbations in peripheral blood and bone marrow hematopoietic cell profiles with features of MPN from an early age (Sandberg, Sutton et al. 2005, Clarke, Volpe et al. 2017). However, the current study shows that if Myb haploinsufficient animals are aged to a point equivalent to old age in humans then a range of myelomonocytic neoplastic phenotypes are seen in the majority of mice, MPN / MDS predominating but with a significant proportion of leukemia, some of which exhibit extramedullary invasion.

### The age-dependent consequences of Myb-insufficiency are largely intrinsic to the HSC

The fact that experimental, constitutive Myb insufficiency only reveals as a hematological disease phenotype in later life, mirroring the association of lower Myb expression as an age-dependent MPN risk factor, raises the question of the reason behind the delay and whether the observed time lag is a consequence of cell intrinsic or extrinsic ageing effects. From studies on both HSC and other tissue stem cells, it has become apparent that the decline in stem cell capability with age, and consequent risk of hematological disease, results from a combination of intrinsic and extrinsic changes, the precise contribution of which is different for individual stem cell types (Mejia-Ramirez and Florian 2020). Cell intrinsic processes affected with age in HSC include DNA damage, cell cycle regulation, epigenetic and transcriptional regulation of gene expression, metabolism, proteostasis, and autophagy, many of which are inter-related. Extrinsic influences on HSC that have been proposed to be age-dependent include the composition and architecture of the immediate bone marrow niche and both local and systemic changes in cytokines and chemokines. Collectively, such changes have the overall effect of increasing proliferation, diminishing self-renewal potential, skewing commitment bias towards myeloid lineages, and limiting the clonal diversity of the population (Sudo, Ema et al. 2000, Pang, Price et al. 2011, Genovese, Schiroli et al. 2014, Jaiswal, Fontanillas et al. 2014).

To distinguish between the possibility of intrinsic versus extrinsic age-related factors impacting on the eventual disease consequences of subnormal Myb levels, a model was established to induce haploinsufficiency in either young or older animals. As for the constitutively haploinsufficient animals, conditional knockout of one *Myb* allele in young mice resulted in signs of MDS / MPN / leukemia after a considerable time lag. Likewise, rendering animals haploinsufficient at one year of age was followed by a considerable time interval before signs of myeloid disease emerged, implying that the ‘clock’ relating to the effects of Myb insufficiency has a major cell intrinsic component.

Given the conclusion that a cell intrinsic defect caused by Myb insufficiency must have a time-dependent effect on HSPC, which is only revealed as a disease phenotype in later life, cell function and molecular differences were compared between WT and *Myb*^+/−^ HSC at both 2 months of age and at 12 months before any signs of hematological changes become apparent. Although peripheral blood cell numbers were not significantly different between WT and *Myb*^+/−^ mice, HSC numbers seemed to be affected by both age and Myb insufficiency, the increase in WT stem cell numbers being in line with what has been previously described (de Haan and Van Zant 1999), while in young *Myb*^+/−^ bone marrow the absolute number of HSC was higher than in wild type mice. Assaying HSC function in terms of long-term reconstitution potential, which puts non-physiological stress on the cells in terms of self-renewal, revealed that both young and old *Myb*^+/−^ HSC are compromised, exhibiting profoundly decreased LT-HSC self-renewal potential and skewing towards myeloid commitment at the expense of lymphoid differentiation.

Transcriptome comparisons enabled a more comprehensive picture of the molecular changes associated with Myb insufficiency and how these might explain the way in which cell intrinsic effects lead to an age-related phenotype. KEGG pathway analysis of the differentially expressed genes suggests that in young adults Myb insufficiency potentially has an impact on DNA replication, metabolism, and especially the proteasome, changes in each of which have been linked with ageing (Mejia-Ramirez and Florian 2020). The degree of effect on the genes associated with these processes is relatively small, being of the order of a 10-20% change in expression compared to normal cells, this degree of difference being reflected in those biological process parameters that could be measured, including proliferation rate, and proteosomal and ribosomal activity.

While similar differences in proteasome-associated genes persist in older animals, a focus on those gene expression differences that are not seen in WT mice as they age reveals a striking increase in the expression of genes encoding proteins associated with ribosome biogenesis, which is reflected as an increase in the rate of protein synthesis. The upturn in ribosome biogenesis would appear to be an indirect age-dependent consequence of Myb insufficiency, which is distinct from the usual age-related changes observed in normal HSC, and it is likely that this results from increased proteosomal activity and the cell’s attempts to maintain proteostasis.

### Myb-insufficiency perturbs HSC proteasome activity in a way that leads to a progressive imbalance in proteostasis

Proteostasis is known to be a key factor in the ageing of tissue stem cells (Schuler, Gebert et al. 2020). The quiescent state that characterises normal adult stem cell maintenance, including in HSC, is characterised by low levels of protein synthesis compared to progenitor cells (Signer, Magee et al. 2014), which serves to reduce the risk of accumulation of misfolded proteins, thereby requiring less effort by the cell with respect to protein clearance. Reflecting this last point, HSC exhibit reduced proteasome activity compared to downstream progenitors (Hidalgo San Jose, Sunshine et al. 2020) and tightly control protein synthesis, at least in part through targets of mTOR (Signer, Qi et al. 2016). In young HSC, the ubiquitin-proteosome system is important for homeostatic regulation of protein degradation, which works synergistically with the regulation of ribosomal biogenesis and protein synthesis.

The increase in proteosome activity could be indicative that there is an imbalance in the requirement of the *Myb*^+/−^ HSC to clear proteins that are potentially mis-folded or unfolded during translation. Accumulation of damaged proteins within the HSC is a hallmark feature of leukemogenesis, and in young *Myb*^+/−^ HSC could potentially create an environment that both hinders their ability to function as a true stem cell but also generate an environment to facilitate the accumulation of other mutations to drive disease. By 12-months of age the Myb haploinsufficient HSC show a highly significant increase in protein synthesis, which in itself should lead to the HSC accumulating a higher number of damaged proteins, thereby perpetuating the phenotype further.

Given its occurrence in young *Myb*^+/−^ mice, the elevated expression of proteasome-associated genes and enhanced proteasome activity could be a direct consequence of the Myb insufficiency. This possibility is supported by the observation that a set of proteasome (*PSM*) genes are bound by MYB in human chronic myeloid leukemia (CML) cells, which are a model of HSPC, and furthermore these genes both respond to *MYB* knockdown in these cells (Lemma, Ledsaak et al. 2021) and are rapidly affected by conditional *Myb* gene deletion in mouse HSC. Most *Psm* genes appear to be influenced by Myb haploinsufficiency, including those encoding the α and β components of the 20S subunit and many of the 19S cap elements that together constitute the 26S proteasome. It is believed that proteasome-associated genes are co-ordinately regulated (Motosugi and Murata 2019), although the mechanisms involved are not fully defined; Myb could be a factor that influences a common regulatory component that underpins this coordination. Analysis of the eight human *PSM*-genes that are homologues of the Myb insufficiency-responsive mouse genes, showed that all are negatively regulated by MYB (three examples are shown in Fig. 6B). Moreover, they are all bound by MYB in their promoter region, showing that they are directly affected by the transcription factor, possibly through some kind of mechanism interfering with their transcriptional initiation.

The link between Myb, proteasome-associated gene regulation, and a propensity for the development of myeloid disease in conditions of Myb insufficiency is further supported by GWAS traits for the key *Psm* genes highlighted in this study. Hence, those *Psm* genes that were identified as being elevated in *Myb*^+/−^ HSC, and in CML cells are both bound by MYB and respond to knockdown, have been linked through GWAS to variations in red cell numbers (*Psma3*, *Psma7*, *Psmb1*, and *Psmg3*), platelet numbers (*Psma7*, *Psmb1*, *Psmd13*, and *Psmg1*), or the occurrence of AML (*Psmb3*).

It is clear that susceptibility to myeloid neoplastic disease can have an inherited component, as is apparent from the occurrence of disease in close relatives of individuals who have already succumbed (Landgren, Goldin et al. 2008, Sud, Chattopadhyay et al. 2018), and that variations at a number of gene loci can be identified as risk factors (Bao, Nandakumar et al. 2020). One of these risk factors involves SNP upstream of the *Myb* gene (Tapper, Jones et al. 2015) while other proteins that influence Myb levels by a direct action on its gene expression, such as Gata2 and Kmt2a (MLL1), are also associated with susceptibility to myeloid disease (Wlodarski, Collin et al. 2017, Katsumura, Mehta et al. 2018, Yin, Xie et al. 2021). Whether an inherited variation in Myb levels is a consequence of genetic differences in the *Myb* locus itself or in genes that encode *Myb* gene regulators, what we have shown here is that the impact on HSPC can be a broad range of proportionately small molecular changes. These accumulated differences result in a combination of constitutive and progressive accumulating shifts in cell behaviour, particularly affecting proteostasis, which could provide the right conditions for clonal expansion, especially in the circumstance when an acquired driver mutation occurs.

## ACKNOWLEDGEMENTS

We are grateful to Paloma García for help in classifying the hematological disease phenotypes and to past members of the Frampton and García groups for their assistance and discussions. We are also grateful to the staff in Genomics Birmingham for next generation sequencing support and the Biomedical Services Unit for animal care.

Computational analysis of the RNA-seq data was performed on the CaStLeS infrastructure at the University of Birmingham (Thompson *et al,* 2019). Part of the bioinformatic analysis was performed on the Saga super computing resources (Project nn9374k) provided by UNINETT Sigma2, the National Infrastructure for High Performance Computing and Data Storage in Norway.

This study was funded by Blood Cancer UK through a Specialist Programme Grant (ref 12030) and a Continuity Grant (ref 17008), and relied upon as MoFlo XDP cell sorter funded through a Wellcome Trust Equipment Grant (084330/Z/07/Z). B.N. is funded through the Cancer Research UK Birmingham Centre award C17422/A25154.

## AUTHOR CONTRIBUTIONS

M.L.C. designed and conducted experiments, analysed data, and wrote the manuscript. L.R.B. and O.S.G. provided RNA-seq and ChIP-seq datasets and their analysis and edited the manuscript. B.N. performed bioinformatic analysis. G.V. collected and processed samples, D.S.W. provided technical support. J.F. designed the study, analysed data, and wrote the manuscript.

## MATERIALS AND METHODS

### Resource availability

#### Lead contact

Further information and requests for resources and reagents should be directed to and will be fulfilled by the lead contact, Mary Clarke.

#### Materials availability

This study did not generate any new unique reagents.

#### Data and code availability

- All RNA-Sequencing data have been deposited <u>in the European Nucleotide Archive (ENA)</u> and are publicly available as of the date of publication. Accession numbers are listed in the key resources table. All data reported in this paper will be shared by the lead contact upon request.
- This paper does not report original code.
- Any additional information required to reanalyze the data reported in this paper is available from the lead contact upon request.

### Experimental model and subject details

#### Mouse models

Animal experiments were carried out in accordance with UK legislation (required that authors identify here the committee approving the experiments). All mice used in this study were controlled using wild type littermates and maintained on a backcrossed C57BL/6J background. Host mice for transplantation assays were male B6.SJL-*Ptprc*^*a*^ / BoyJ. For conditional deletion of *Myb*, a cohort of male *Myb*^+/*F*^:*CreER*^*T2*^ (Emambokus, Vegiopoulos et al. 2003) mice together with *Myb*^+/+^:*CreER*^*T2*^ mice were randomly assigned to groups for deletion at either 2- or 12-months of age.

### Method details

#### Ageing mouse model

All mice were genotyped by ear clip sampling and genotyped by PCR using DreamTaq polymerase (ThermoFisher, Waltham MA). The primers were: (i) For the intron 6 loxP site, ATCTGAAGAAAATGAATTGA and GCATCAGCTCGATGATAAGCA giving products of 233 bp and 281 bp from the wild-type and loxP-modified loci; ii) Detection of the mTmG transgene using primers against eGFP, GTAAACGGCCACAAGTTCAG and GTCTTGTAGTTGCCGTCGTC, and Cre, TCGATACAACGAGTGATGAG and TTCGGCTATACGTAACAGGG giving products of 260bp and 500bp respectively; (iii) For the *Myb* null allele, CCATGCGTCGCAAGGTGGAAC and TGGCCGCTTTTCTGGATTCATC giving an amplification product of 295 bp; iv) for the vWF transgene CCTCTCTGGACGGTGAGAAC and GTCTTGTAGTTGCCGTCGTC giving a product of 312bp for the transgene.

A cohort of *Myb*^+/−^ mice together with littermate WT controls were bled monthly by the tail vein into 30 μl of acid citrate dextrose (ACD: 6.8 mM citric acid; 11.2 mM trisodium citrate; 24 mM glucose) anti-coagulant solution and analysed on an automated ABX Pentra 60 blood counter (Horiba, Northampton UK). Blood values were calculated according to the dilution ratio of blood:ACD. All mice were aged up until 22 months of age, unless they were culled due to initiation of disease (determined by monthly blood values, body weight and condition) and culled by cervical dislocation and femurs, tibias, spleen, liver and lungs dissected for analysis. The spleen and liver were weighed before all organs and one tibia fixed in 4% paraformaldehyde. Tibias were decalcified prior to embedding in paraffin wax and sectioning. Tissue sections were stained with Hematoxylin:Eosin before analysis by microscopy.

Conditional deletion of *Myb* gene alleles was induced by intraperitoneal injection of 75mg/kg Tamoxifen (Sigma, St Louis MO) dissolved in corn oil. Deletion of floxed *Myb* gene alleles was determined by preparing DNA from peripheral blood cells and performing PCR to quantify the loxP-flanked region of *Myb* (exon 5) (5’-ATCTGTTCCCCAGACGCTTG-3’ and 5’-ATGACGCTGACTCTTTCCGT-3’) normalised against *Itga2b* as a control gene (5’-AACTGTTTGTGGACGGAGTCACTG-3’ and 5’-GATTCAGCCTTTCAGCAGCACTAC-3’).

### Flow cytometry and cell sorting

Single cell suspensions of bone marrow cells harvested from the tibia and femurs of mice were red cell-lysed in Ammonium-Chloride-Potassium (ACK) buffer, counted, resuspended in PBS-10%FBS and stained for surface antigens. All antibodies were purchased from eBioscience (San Diego CA) with the exception of Sca1 and CD48 that were purchased from BD Biosciences (Franklin Lakes, NJ). Lineage-positive (Lin^+^) cells were defined using monoclonal antibodies against CD5, CD8a, CD11b, B220, Gr1, Ter119. The pool of HSC and multipotent progenitors were defined as Kit^+^Sca1^+^Lin^−^ (‘KSL’), which were further subdivided based on their expression of the SLAM markers CD48 and CD150. Analysis of differentiated cells used antibodies against Gr1, CD11b, B220, CD43, CD4, CD8a, CD41. Cells were analysed on a CyAn ADP (Beckman Coulter, Brea CA) or isolated to 95% or greater purity on a MoFlo XDP high-speed cell sorter (Beckman Coulter, Brea CA), data was analysed using Summit 4.3 (Dako Cytomation).

### Engraftment potential of stem cells

To assess the engraftment potential of HSC from WT and *Myb*^+/−^ mice, sorted KSL cells were transplanted by intravenous injection into the tail vein of B6.SJL (*Ptprc*^*a*^) host animals that had been lethally irradiated (900 Gy). Test donor mice carried the *vWf-tomato* transgene for assessment of donor platelet differentiation in the periphery. Test donor cells (CD45.2/CD45.2) were injected together with red cell-lysed whole bone marrow from C57BL6:B6.SJL F1 generation (CD45.1/CD45.2) mice (5 × 10^5^ cells), the latter acting as both a reference and to provide stable haematopoiesis. Mice were bled from the tail vein once a month into ACD solution to determine the ratio of engraftment between test and reference donor using immunofluorescent flow cytometric analysis to quantify CD45.1 and CD45.2 antigens.

### RNA-Seq analysis

RNA-Seq was performed using the NEBNext® Single Low Input RNA Library Prep and sequencing. The sequencing platform was NextSeq 500 (Illumina), using a v2.5 150 cycles Mid output (75 paired end) Flowcell. Sequencing reads were mapped to GRCm38 (mm10) mouse genome with STAR aligner (v2.5.2b) (Dobin, Davis et al. 2013). Reads mapping to genes were counted by the same software. Normalisation of read counts and differential expression analysis were performed with DESeq2 (v.1.26.0) R Bioconductor package (Love, Huber et al. 2014). Gene set enrichment analysis was done using GAGE (v.2.36.0) R Bioconductor package with Gene Ontology databases (Luo, Friedman et al. 2009).

### Proteosome activity assay

WT and *Myb*^+/−^ bone marrow cells stained for the KSL markers were incubated with 5μM Me4bodipyFL-Ahx3Leu3VS probe (R&D Systems, Minneapolis MN) for 1 hour at 37°C either with or without prior incubation for 1 hour at 37°C with 10μM of the proteosome inhibitor MG-132 (Millipore Sigma, St Louis MO). Following incubation, cells were fixed in 1.6% paraformaldehyde and analysed by flow cytometry.

### Protein synthesis assay

Protein synthesis was assayed by labelling WT and *Myb*^+/−^ bone marrow cells stained for the KSL markers with O-propargyl-puromycin (OPP) followed by Click-iT technology (Invitrogen, Waltham MA) as per the manufacturers protocol and assayed by flow cytometry.

### Proliferation assay

WT and *Myb*^+/−^ bone marrow cells were stained for the KSL markers followed by fixation in 1.5% paraformaldehyde and permeabilization in ice cold acetone. Cells were then stained with eFluor 450-conjugated anti-Ki67 antibody (clone SolA15, ThermoFisher, Waltham MA) and then analysed by flow cytometry.

### Autophagy assay

Bone marrow cells from WT and *Myb*^+/−^ mice were stained for the KSL surface markers followed by fixation in 3.7% paraformaldehyde and permeabilization in ice cold acetone. Cells were then stained for 30 minutes with an antibody against LC3B (clone 1251A) conjugated to APC, or an anti-rabbit IgG APC control (Novus Biologicals, Littleton CO) at a 1:100 dilution.

### Gene expression analysis

RNA was extracted from sorted KSL cells using TRIzol and converted to cDNA using M-MLV reverse transcriptase (Promega, Madison WI). Quantitative PCR was performed using Taqman (Applied Biosystems, Waltham, MA) for B2m (Mm00437762_m1), Psmb3 (Mm04210356) and Rps8 (Mm01187191_g1).

### Bioinformatic comparison of mouse HSC transcriptome data with genome-wide MYB chromatin-binding data from human haematopoietic progenitor cells

In order to compare the affected genes from *Myb*^+/−^ HSC with their human counterpart, we took advantage of the recently published MYB ChIP-seq data from the K562 cell line (Lemma, Ledsaak et al. 2021). We focussed on direct target genes of MYB, defined as genes that respond to MYB knockdown and rescue (Fuglerud, Lemma et al. 2017) and being occupied by one or more of MYB ChIP-seq peaks (Lemma, Ledsaak et al. 2021). Target genes of MYB with human gene IDs were converted to their corresponding mouse genes through a combination of two resources; (1) the mouse genome database (MGD) accessible through Mouse Genome Informatics (MGI) (Accessed on 18^th^ of May, 2021) (Bult, Blake et al. 2019), and (2) the *Homologene* R package (version 1.4.68.19.3.27) (Mancarci & French, 2019), resulting in n=523 genes. This gene set was compared with Myb insufficiency-responsive genes assayed by RNA-seq of HSC from 2-month-old WT and *Myb*^+/−^ mice, the latter restricted to those Myb insufficiency-responsive genes with significant DE between the two conditions and meeting stringent filtering criteria, where absolute value of log2FC ≥ 0.5 and *P*-value < 0.05, resulting in n=662 significantly DE genes. Finally, the intersection between the direct MYB target genes was converted to mouse gene IDs (n=523) and the significantly DE Myb insufficiency-responsive genes (n=662) were determined.

Pathway and functional term enrichment analysis was performed on the list of genes from the intersection step above using the *msigdbr* function from *msigdbr* R package (version 7.2.1) to connect to the molecular signature database (MsigDB) and retrieve the curated C2 set for mouse. Then, the *enricher* function from *clusterProfiler* R package version 3.12.0) (Yu et al., 2012) was employed to identify gene set enrichment in the intersection list with that of the C2 curated list of genes from MsigDB (Dolgalev, 2020), considering only the top significant enriched terms ranked according to Benjamini and Hochberg adjusted P-value < 0.05.

### Quantification and statistical analysis

For all experiments, significant changes were calculated using a two-tailed Student’s t-test with a significance threshold of P<0.05 with the exception of the Kaplan-Meier survival analysis that was calculated using the LogRank test with significance threshold of <0.05. Significance values can be found in the figure legends, where ‘n’ is the number of animals in the cohort analysed. All bar graph plots depict the mean result with error bars representing the standard error of the mean (SEM). In bone marrow transplantation assays, ‘n’ represents the number of recipient mice analysed.

Quantitative PCR was performed using experimental triplicates and repeated using multiple samples therefore n is the number of biological replicates used. Data was analysed using both 2-ΔΔCT and 2-ΔCT methods. Average relative expressions were plotted in Excel (Microsoft, Redmond WA) with error bars representing the SEM.

### Additional resources

OPP assay link:https://www.thermofisher.com/document-connect/document-connect.html?url=https://%3A%2F%2Fassets.thermofisher.com%2FTFS-Assets%2FLSG%2Fmanuals%2F10456_Click_iT_PlusOPP_Protein_SynthesisAssayKits_PI.pdf&title=Q2xpY2staVQgUGx1cyBPUFAgUHJvdGVpbiBTeW50aGVzaXMgQXNzYXkgS2l0cw==

## SUPPLEMENTAL INFORMATION

**Figure S1.**
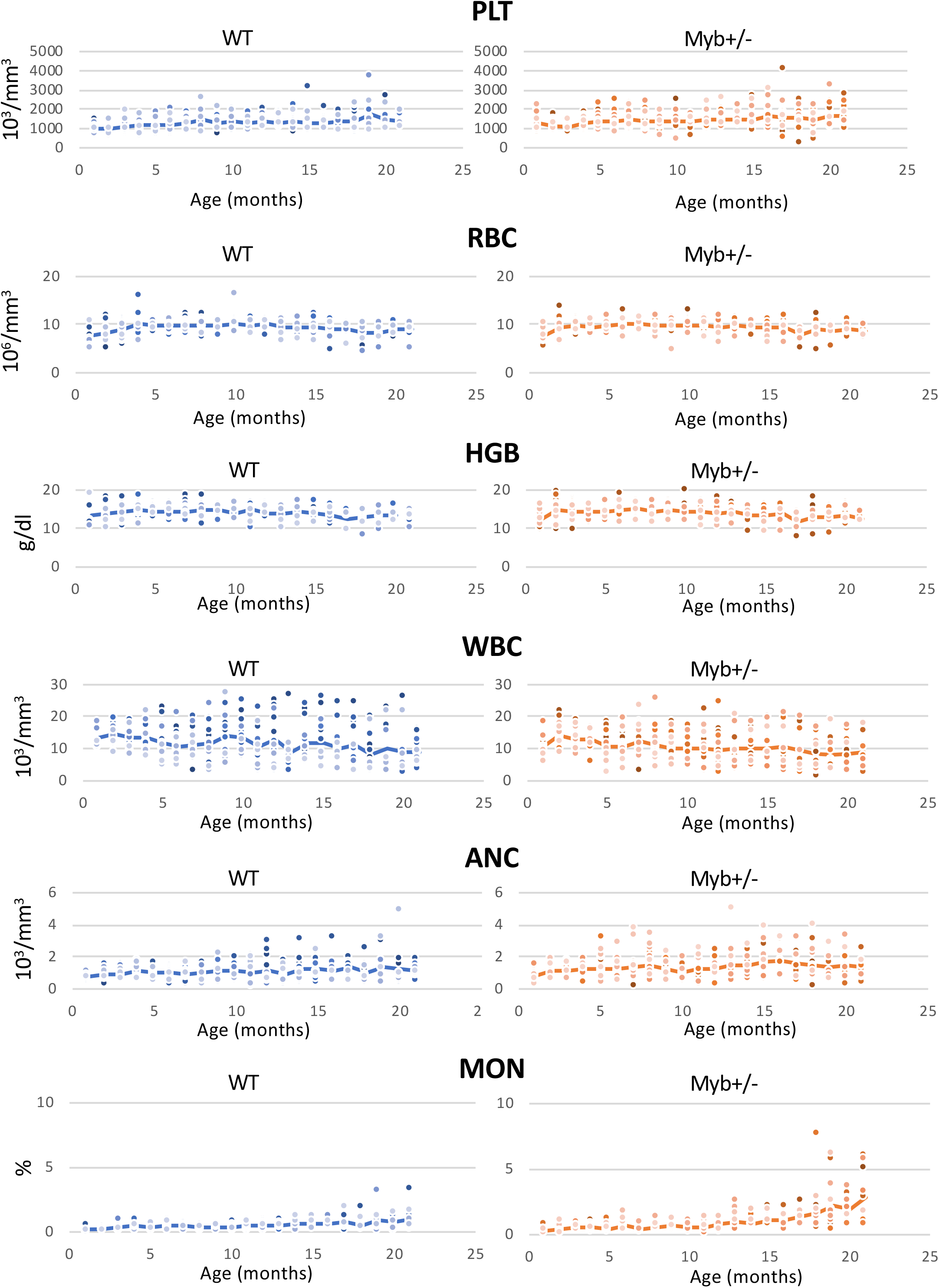
Peripheral blood values in aging cohorts of WT and *Myb*^+/−^ mice. Peripheral blood counts showing individual mice from aged wild type (WT) (n=17) and *Myb*^+/−^ (n=19) cohorts over 22 months. PLT (platelet), RBC (red blood cell), HGB (Hemoglobin), WBC (white blood cell), ANC (absolute neutrophil count), MON (monocyte).

**Figure S2.**
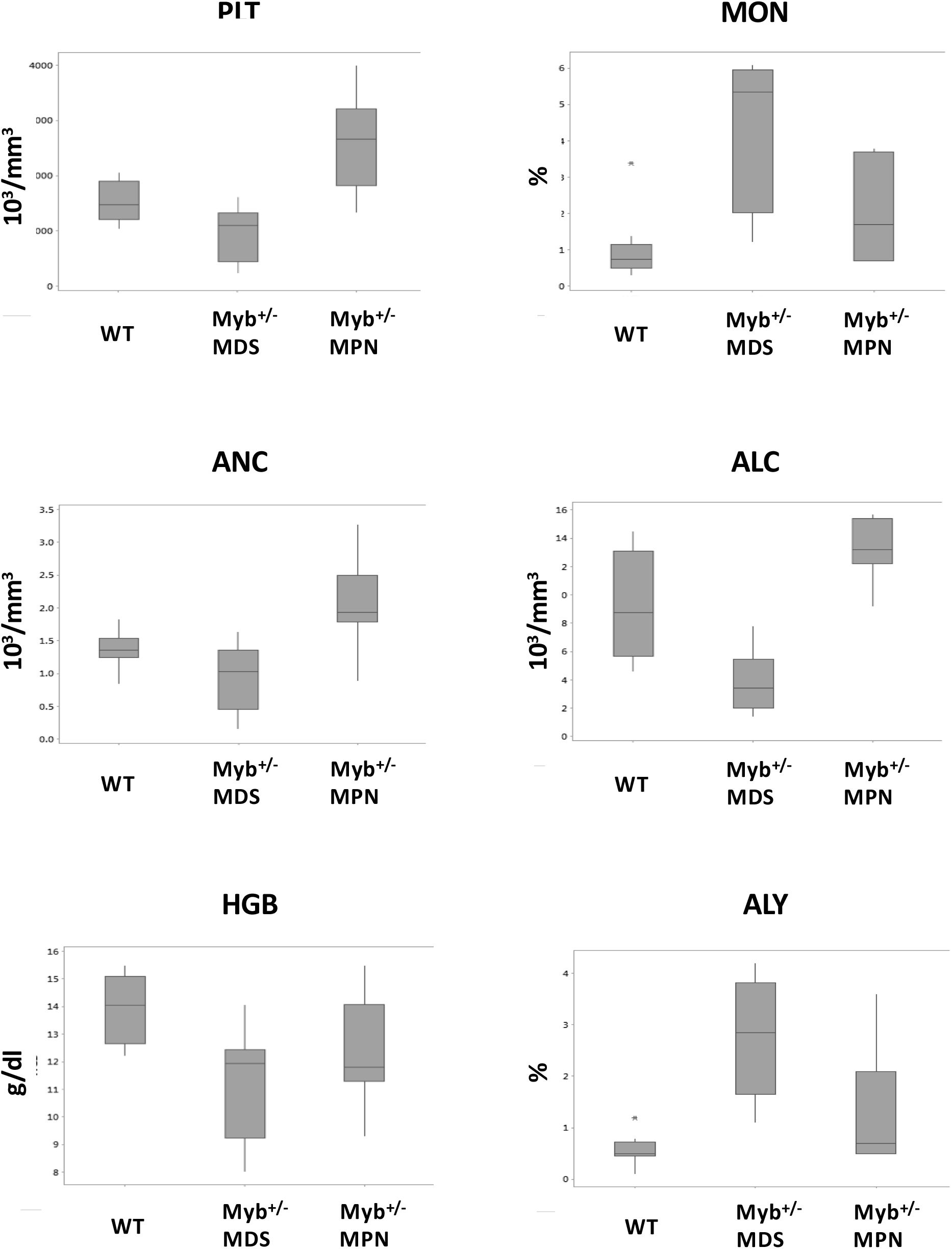
Peripheral blood values in aged WT and *Myb*^+/−^ mice at time of sacrifice. Peripheral blood counts at date of sacrifice of aged WT and *Myb*^+/−^ mice categorised into the disease phenotype of MDS and MPN. PLT (platelet), MON (monocyte), ANC (absolute neutrophil count), ALC (absolute lymphocyte count), HGB (hemoglobin), ALY (atypical lymphocytes).

**Figure S3.**
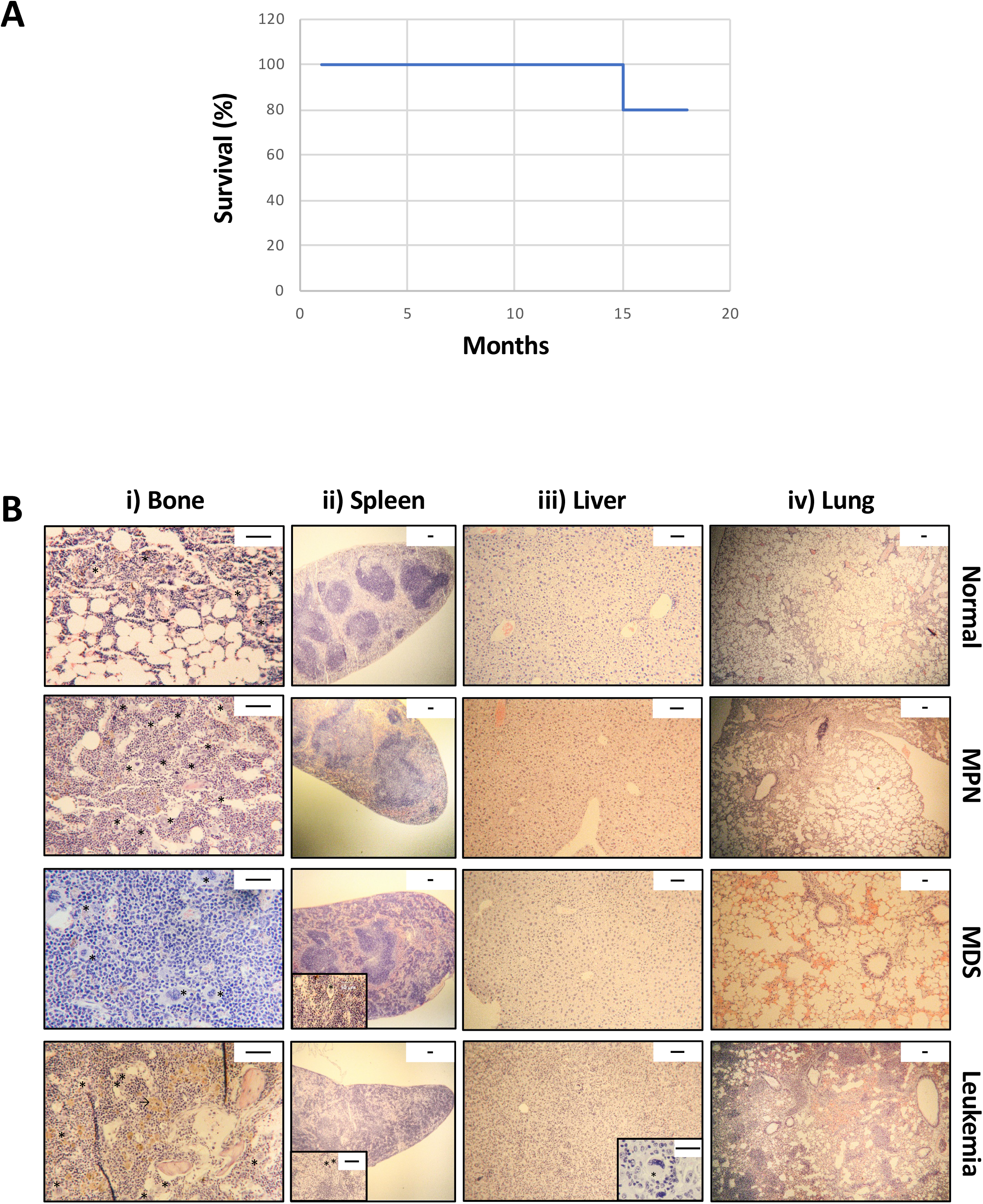
Induction of Myb deletion with Tamoxifen. A) Survival curve of Myb+/− mice (n=5) injected with Tamoxifen at 2 months of age. B) Representative histology sections of tibial bone, spleen, liver and lung from *Myb*^+/+^:*CreER*^*T2*^:*mTmG* and *Myb*^+/*F*^:*CreER*^*T2*^:*mTmG* mice that were induced to delete Myb at 2- or 12-months of age. Images show an example representing each disease phenotype. i) Bone sections with megakaryocytes (*). Scale bar = 50μm. ii) Spleen sections showing the presence of extramedullary megakaryocytes (inset * scale bar = 50μm). Scale bar = 100μm. iii) Liver sections with infiltrating megakaryocytes (inset * scale bar = 50μm). Scale bar = 100μm. iv) Sections of lung showing involvement of myeloid cells. Scale bar = 100μm.

**Figure S4.**
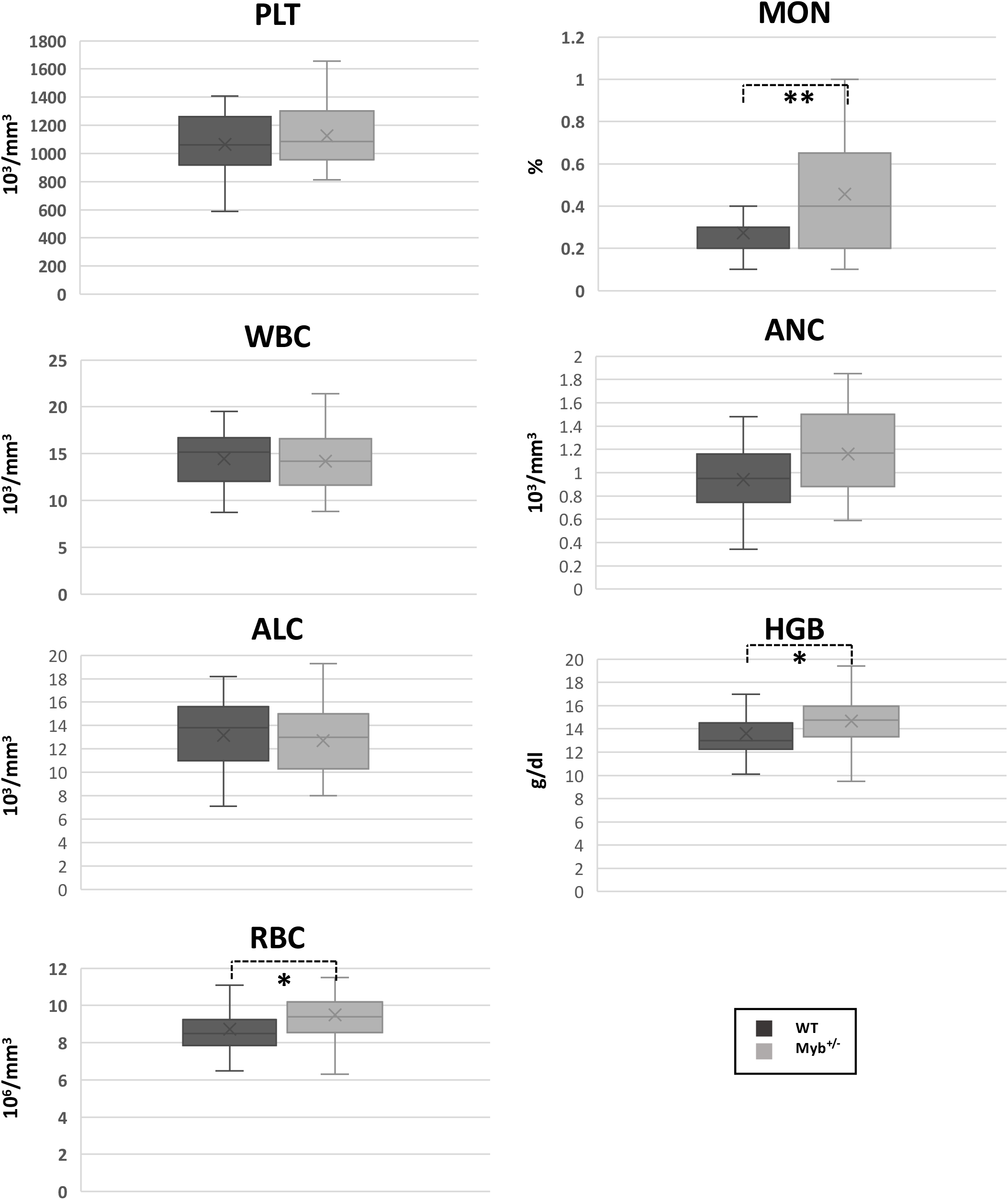
Peripheral blood values of WT and *Myb*^+/−^ mouse cohorts at 2 months of age. Peripheral blood counts of a cohort of 2-month old wild type (WT) (n=17) and *Myb*^+/−^ (n=19) mice. PLT (platelet), MON (monocyte), WBC (white blood cell), ANC (absolute neutrophil count), ALC (absolute lymphocyte count), HGB (hemoglobin), RBC (red blood cell). p^mon^ = 0.005, p^HGB^= 0.040, p^RGB^= 0.031.

**Figure S5.**
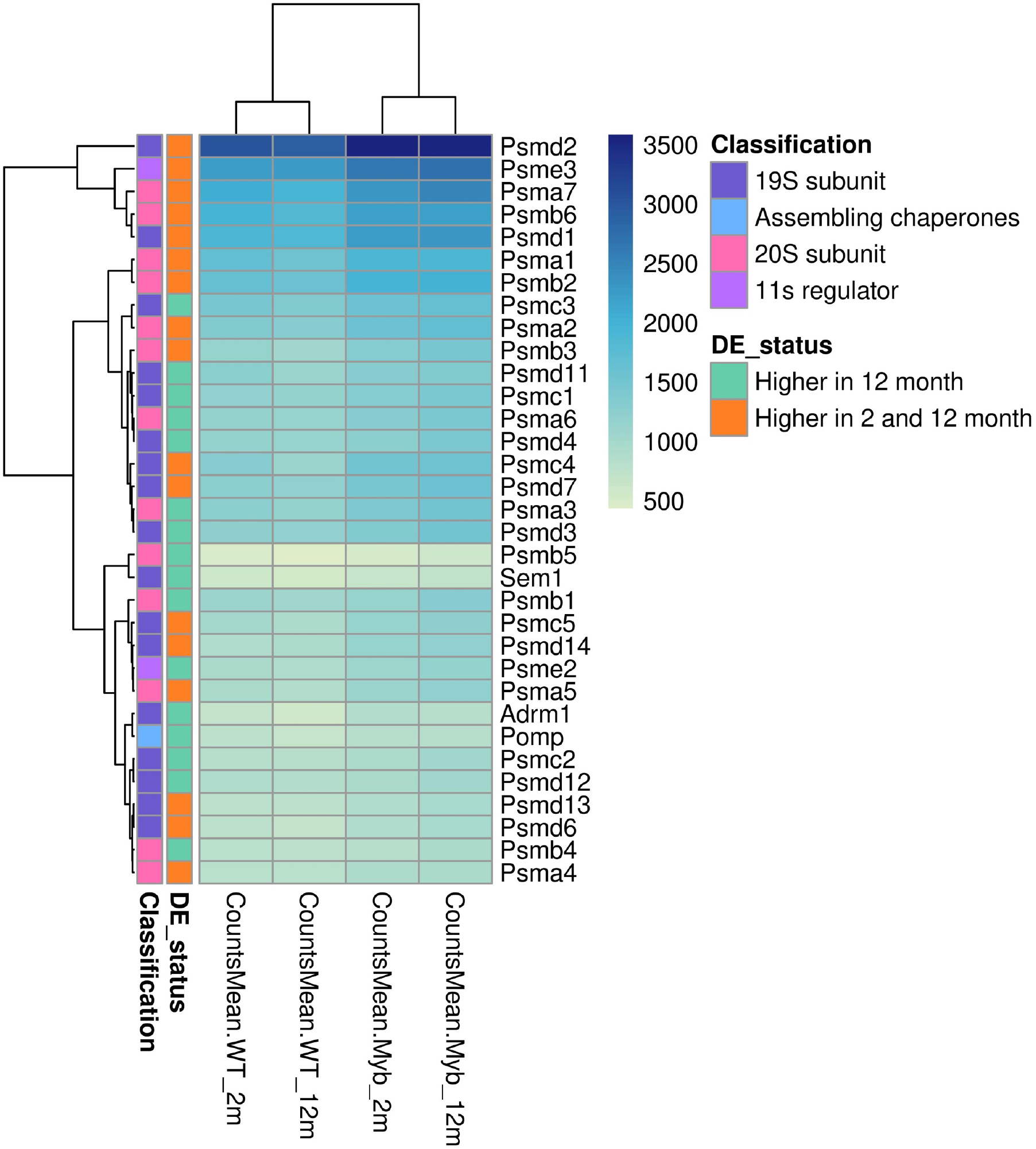
Proteosome component gene expression comparison at 2- and 12-months for WT versus *Myb*^+/−^ HSC. Expression heatmap of genes that belong to the (1) 19S subunit, (2) the 20S subunit, (3) the 11s regulator, and (4) assembling chaperon categories, which showed higher gene expression at either 12-months only or at both 2- and 12-months. The heatmap was generated by hierarchical clustering of the RNA-seq data for these genes using the euclidean distance and complete agglomeration method. The classification of the genes into the different proteasome regulatory particles is displayed as an annotation next to each gene. Similarly, the status of differential gene expression is displayed as an annotation next to each gene. The heatmap was generated using the pheatmap R package version 1.0.12 (Kolde, 2019).

**Figure S6.**
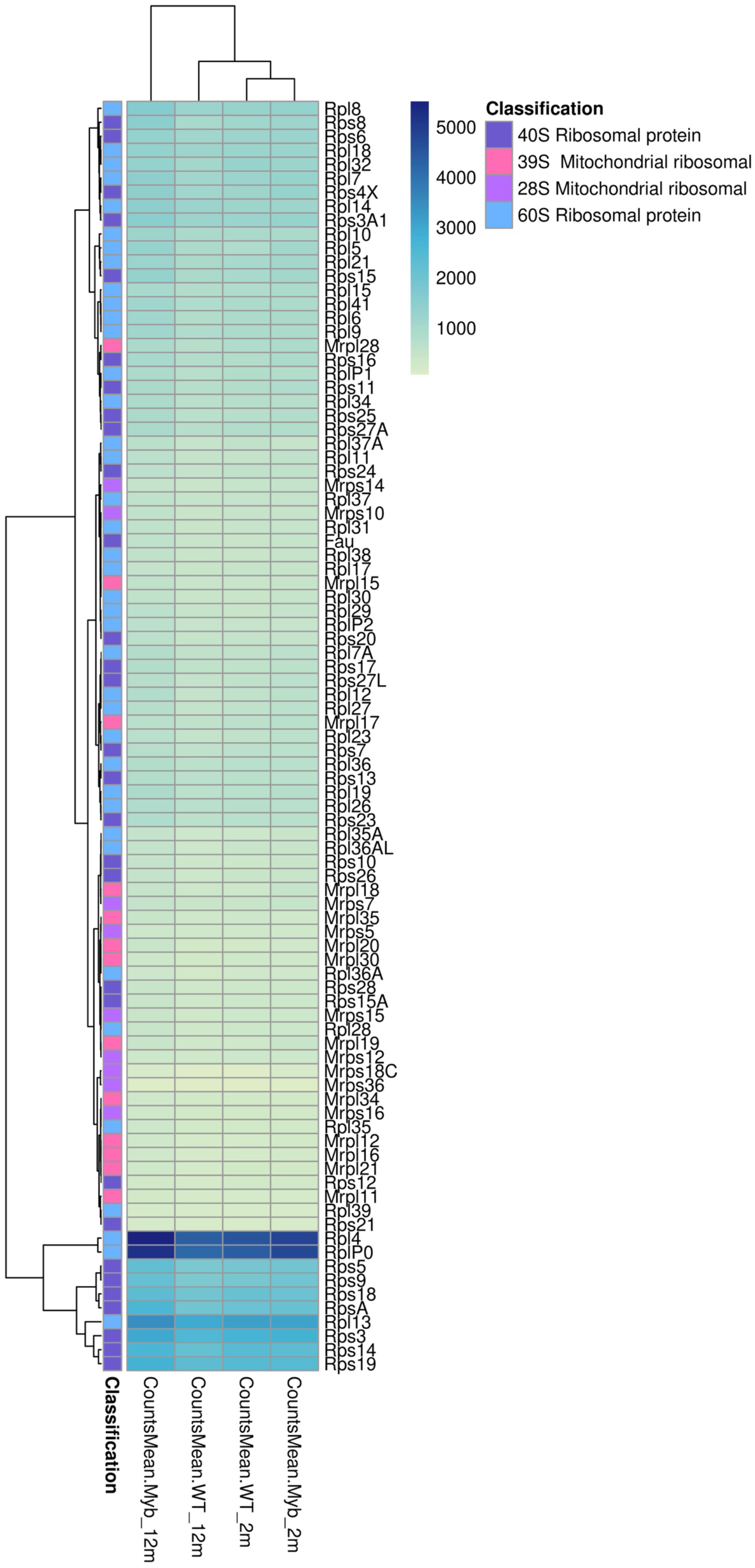
Ribosome component gene expression comparison at 2- and 12-months for WT versus *Myb*^+/−^ HSC. Expression heatmap of genes that belong to the (1) 28S mitochondrial ribosomal, (2) 39S mitochondrial ribosomal, (3) 40S ribosomal protein, and (4) 60S ribosomal protein categories, which exhibited DE at 12-months. The heatmap was generated as described in Supplementary Figure S5.

**Figure S7.**
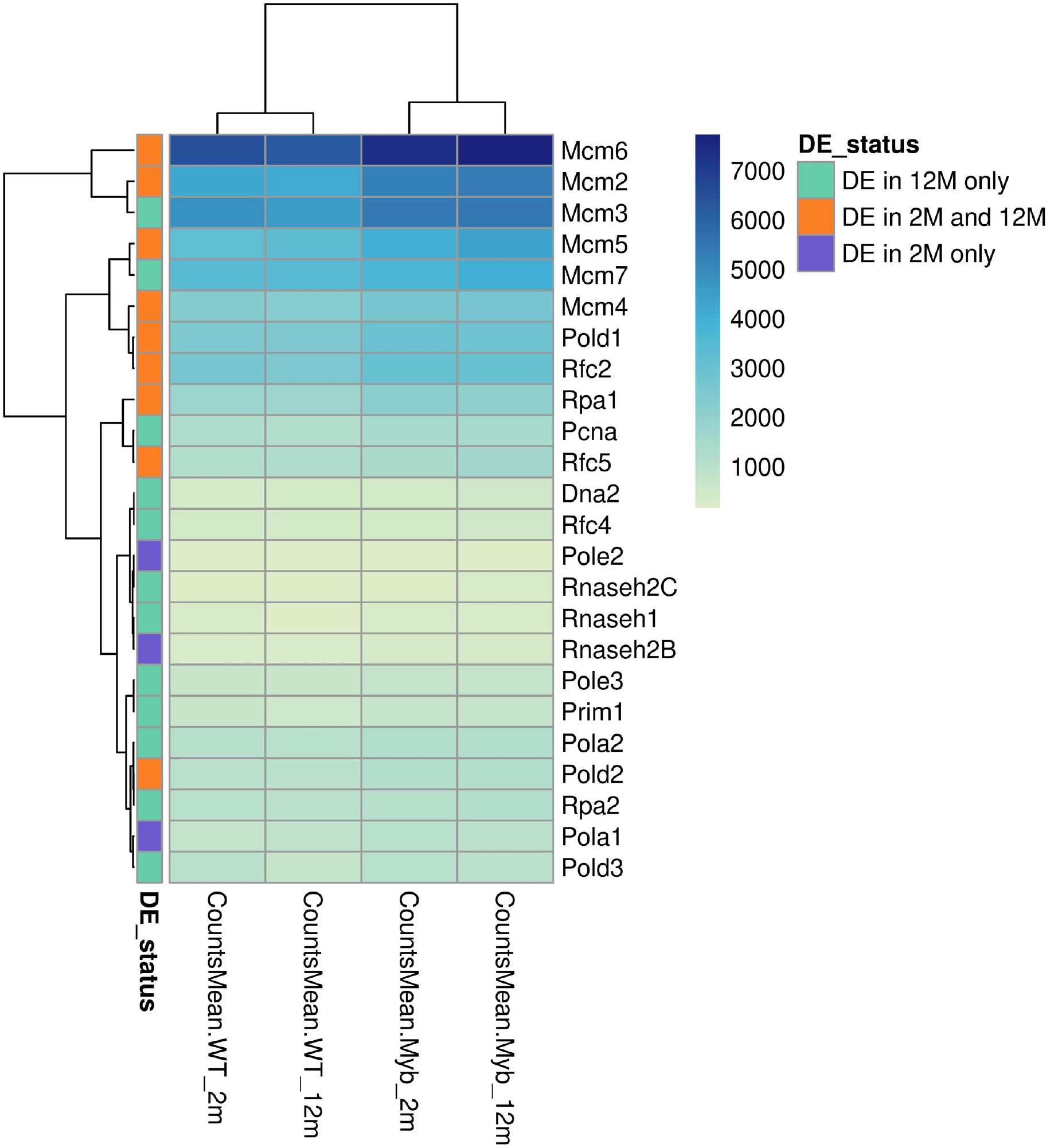
DNA replication-associated gene expression comparison at 2- and 12-months for WT versus *Myb*^+/−^ HSC. Expression heatmap of proliferation genes that showed DE at (1) 2-months only, (2) 12-months only, and (3) both 2- and 12-months. The heatmap was generated as described in Supplementary Figure S5.

**Figure S8.**
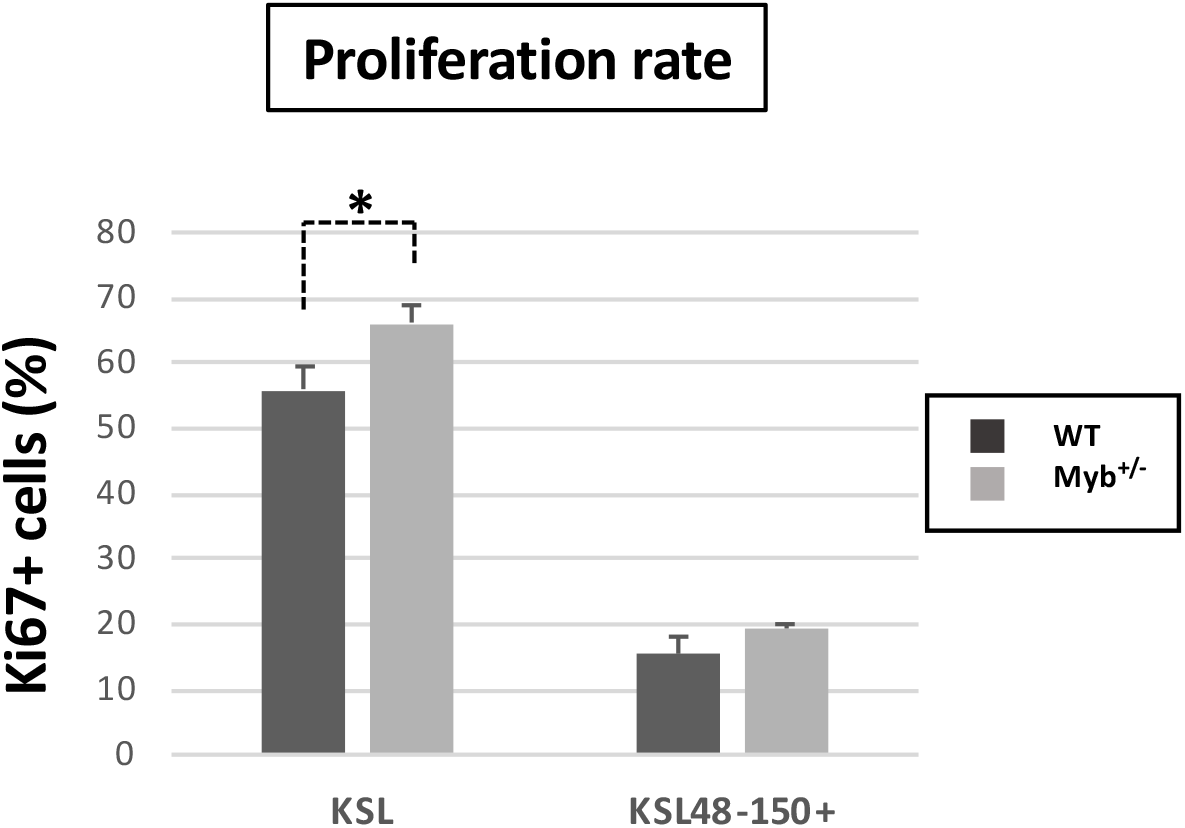
Proliferation rate of WT versus *Myb*^+/−^ HSC from 2-month-old mice. Bone marrow from WT and *Myb*^+/−^ mice (n=5) were fixed and permeabilised for intracellular anti-Ki67 staining. KSL HSC and LT-HSC (KSL48-150+) cells were gated for analysis of Ki67. Data represents the average percentage of Ki67^+^ cells with SEM. For KSL HSC P= 0.024.

